# PlantCAD2: a DNA foundation model for interpreting genomes across flowering plants

**DOI:** 10.1101/2025.08.27.672609

**Authors:** Jingjing Zhai, Aaron Gokaslan, Sheng-Kai Hsu, Szu-Ping Chen, Zong-Yan Liu, Edgar Marroquin, Eric Czech, Betsy Cannon, Ana Berthel, M. Cinta Romay, Matt Pennell, Volodymyr Kuleshov, Edward S. Buckler

## Abstract

Understanding how DNA sequence encodes biological function remains a fundamental challenge in biology. Flowering plants (angiosperms), the dominant terrestrial clade, exhibit maximal biochemical complexity, extraordinary species diversity (over 100,000 species), relatively recent origins (∼160 million years), ∼200-fold variation in genome size and relative compact coding regions compared with other eukaryotes. These features present both a unique challenge and opportunity for pre-training DNA language models to understand plant-specific evolutionary conservation, regulatory architectures and genomic functions. Here, we introduce PlantCAD2, an extended context, plant-specific DNA language model with single-nucleotide resolution, pre-trained on 65 angiosperm genomes, together with a series of public benchmarks for evaluation. Comprehensive zero-shot testing shows that PlantCAD2 (676 million parameters) efficiently captures evolutionary conservation, surpassing the 7-billion-parameter Evo2 model in 10 of 12 tasks. With parameter-efficient fine-tuning, PlantCAD2 also outperforms the 1-billion-parameter AgroNT across seven cross-species tasks including chromatin accessible region, gene expression and protein translation. Moreover, its 8,192bp context window substantially improves accessible chromatin prediction in large genomes such as maize (AUPRC increasing from 0.587 to 0.711), underscoring the importance of long-range context for modeling distal regulation. Together, these results establish PlantCAD2 as a powerful, efficient, and versatile foundation model for plant genomics, enabling accurate genome annotation across diverse species.

## Introduction

Deciphering how DNA sequence encodes molecular functions, phenotypes and fitness remains a fundamental goal in biology. The rapid decline in sequencing costs has enabled large-scale initiatives such as the Darwin Tree of Life project ^1^, the Earth BioGenome Project ^2^, the Vertebrate Genomes Project ^3^, and the 10KP Plant Genome Project ^4^, which collectively aim to sequence tens of thousands of species across the tree of life, with plants alone contributing over a thousand assembled genomes ^5^. While genomic data accumulates exponentially, functional annotations lag far behind, particularly in plants where labeled data exists for only a few model species and crops ^6^. Flowering plants (angiosperms) ^7^, the dominant terrestrial clade ^8^, underpin global food security and ecosystem function. They exhibit maximal biochemical complexity, extraordinary species diversity (over 300,000 species), wide genome size variation, and relatively compact coding regions compared with other eukaryotes—features that present both a unique challenge and an opportunity for computational approaches. This widening gap between available sequences and functional knowledge highlights the critical need for models that can learn from raw sequences alone and transfer knowledge across plant species.

Foundation models ^9^, large neural networks pre-trained on vast collections of unlabeled data using self-supervised objectives have emerged as a powerful approach to this challenge. Rather than requiring large, labeled datasets for each task, these models learn general representations that can be adapted to specific applications with minimal supervision. This paradigm has achieved significant success in protein science, where models such as ESM ^10–12^, ProtTrans ^13^, and ProBERT ^14^ have demonstrated strong performance in predicting protein function ^15^, structure ^16^, and variant effect ^17^.

In contrast, genomic LMs are still rapidly evolving, with recent developments spanning DNA ^18–27^, RNA ^28–30^ and transcriptomes ^31–33^. Early work such as the DNABERT^20^ pre-trained BERT ^34^ model on the human genome showed improved regulatory element prediction. Multi-species pre-training soon proved critical for learning evolutionary conservation^21,22,25,35^, a principle first demonstrated in plants by GPN^22^, which was pre-trained on eight Brassicales genomes and achieved strong variant effect prediction. Broader multi-species models followed, including Nucleotide Transformer ^25^ (850 genomes excluding plants) and Evo ^18^ (prokaryotic and phage genomes). Within plants, AgroNT ^36^ extended pre-training to 48 genomes with longer context windows but sacrificed single-nucleotide resolution through non-overlapping k-mer tokenization. We previously developed PlantCaduceus ^27^ (PlantCAD), a DNA LM pre-trained on 16 divergent angiosperm genomes. It uses the Caduceus ^26^ architecture, a Mamba-based ^37^ design that efficiently models both DNA strands simultaneously. PlantCAD achieved up to a 7-fold improvement over the next-best model in cross-species gene annotation tasks and variant effect prediction tasks. However, its context window of 512 base pairs restricts its ability to model biological processes that depend on distal regulatory interactions ^38^. Many regulatory elements influence gene expression over tens to hundreds of kilobases and are key contributors to phenotypic variation ^39–41^, yet remain challenging to capture with short-context models.

Most recently, Evo2 ^19^ has scaled to 40 billion parameters trained across genomes spanning the entire tree of life. While such generalist models represent impressive engineering achievements, deploying a 40 billion parameter model for genome-wide annotation across more than 300,000 angiosperm species is prohibitively expensive, placing it beyond the reach of most plant genomics laboratories. More fundamentally, pre-training across all domains of life may dilute the lineage-specific regulatory signals critical for any single clade.

In this paper, we introduce PlantCAD2, an improved DNA LM tailored to angiosperm genomes. PlantCAD2 is pre-trained using a masked language modeling objective on 65 curated flowering plant genomes. PlantCAD2 is built on the efficient Mamba2 architecture ^42^, which scales linearly with sequence length instead of quadratically as in transformers ^43^. It supports 8,192-bp input windows and reverse-complement equivariance, allowing the model to capture extended-context, strand-invariant regulatory features. To reduce pretraining bias, we applied sampling strategies that both down-weight repetitive sequences and emphasize coding and regulatory regions. Subsequently, we first evaluated PlantCAD2 on 12 comprehensive benchmarks using a zero-shot strategy, demonstrating its efficiency and capacity to capture evolutionary conservation (**Table 1**). We then fine-tuned the model on seven functional genomics tasks including chromatin accessibility, gene expression, and protein abundance to further demonstrate its state-of-the-art cross-species predictive ability (**Table 2**). Together, these results highlight PlantCAD2’s ability to generalize across species and tasks, and to serve as a versatile foundation model for plant genome interpretation.

**Table 1.**
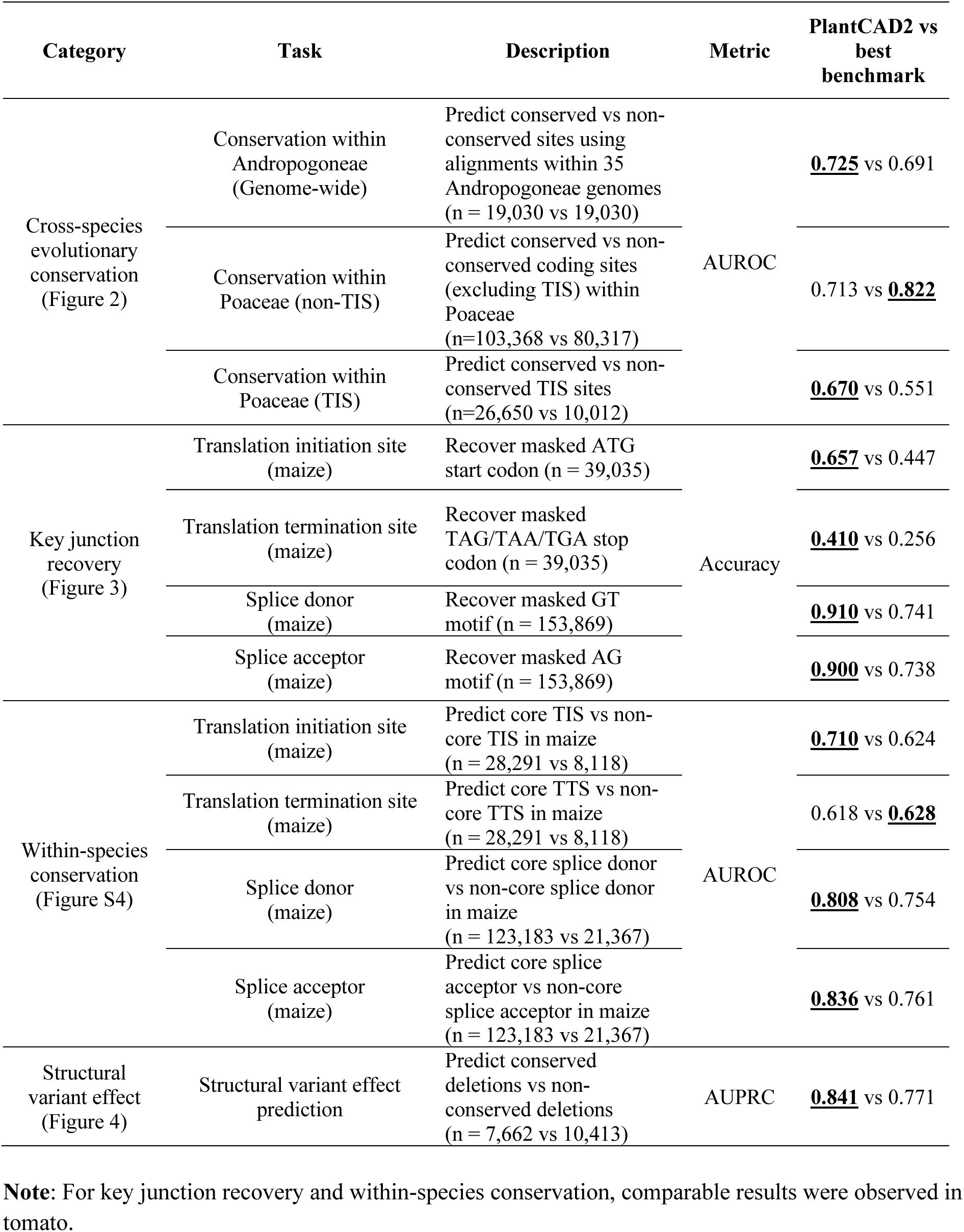
Zero-shot evaluation summary compared with the best-performing benchmark models. For each task, the bold and underscored value indicates the highest score.

**Table 2.**
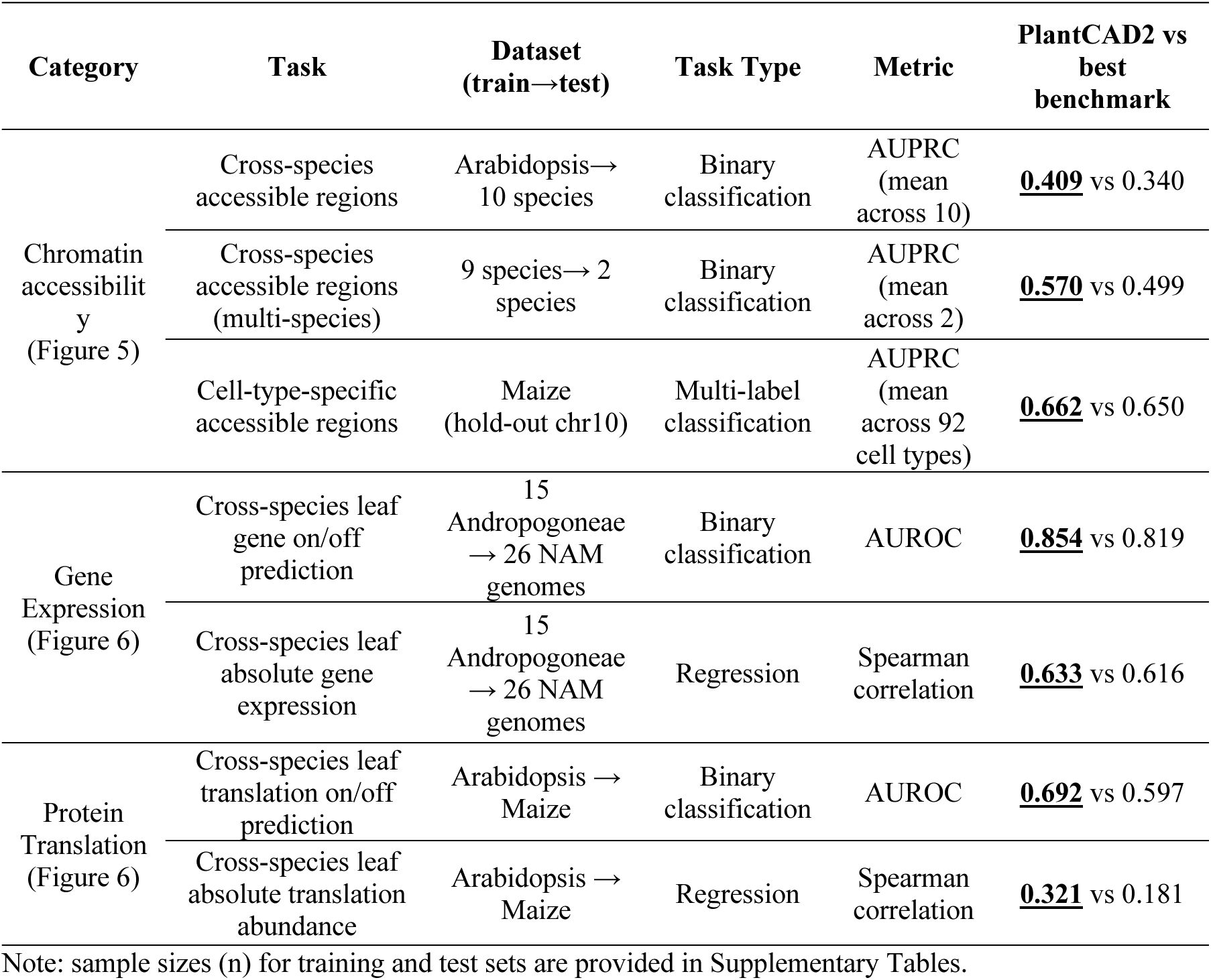
Fine-tuning evaluation summary compared with the best-performing benchmark models. For each task, the bold and underscored value indicates the highest score.

| Category | Task | Dataset<br>(train→test) | Task Type | Metric | PlantCAD2 vs<br>best<br>benchmark |
| --- | --- | --- | --- | --- | --- |
| Chromatin<br>accessibilit<br>y<br>(Figure 5) | Cross-species<br>accessible regions | Arabidopsis→<br>10 species | Binary<br>classification | AUPRC<br>(mean<br>across 10) | **<u>0.409</u>** vs 0.340 |
|  | Cross-species<br>accessible regions<br>(multi-species) | 9 species→ 2<br>species | Binary<br>classification | AUPRC<br>(mean<br>across 2) | **<u>0.570</u>** vs 0.499 |
|  | Cell-type-specific<br>accessible regions | Maize<br>(hold-out chr10) | Multi-label<br>classification | AUPRC<br>(mean<br>across 92<br>cell types) | **<u>0.662</u>** vs 0.650 |
| Gene<br>Expression<br>(Figure 6) | Cross-species leaf<br>gene on/off<br>prediction | 15<br>Andropogoneae<br>→ 26 NAM<br>genomes | Binary<br>classification | AUROC | **<u>0.854</u>** vs 0.819 |
|  | Cross-species leaf<br>absolute gene<br>expression | 15<br>Andropogoneae<br>→ 26 NAM<br>genomes | Regression | Spearman<br>correlation | **<u>0.633</u>** vs 0.616 |
| Protein<br>Translation<br>(Figure 6) | Cross-species leaf<br>translation on/off<br>prediction | Arabidopsis →<br>Maize | Binary<br>classification | AUROC | **<u>0.692</u>** vs 0.597 |
|  | Cross-species leaf<br>absolute translation<br>abundance | Arabidopsis →<br>Maize | Regression | Spearman<br>correlation | **<u>0.321</u>** vs 0.181 |

Note: sample sizes (n) for training and test sets are provided in Supplementary Tables.

## Results

### PlantCAD2: an extended-context DNA language model for angiosperms

PlantCAD2 builds on the original PlantCAD ^27^ DNA language model, preserving its single-nucleotide tokenization and masked language modeling objective, while introducing four major improvements: architectural efficiency, context length, parameter scale, and phylogenetic breadth (**Figure 1A**). First, PlantCAD2 retains the Caduceus ^26^ architecture with its bidirectional, reverse-complement-equivariant design, but replaces the original Mamba ^37^ blocks with Mamba2 blocks ^42^. Mamba2 has substantial advantages over Mamba1, leveraging structured state space duality for more efficient parallel training and simplifying recurrence computations to reduce memory usage (see Methods). Compared to traditional transformer architectures ^43,44^, PlantCAD2 model architecture shows a much slower increase in inference time than modernBERT models under the same input and output dimensions (**Figure S1A**), due to the inherent efficiency of state space models in handling long sequences ^37,45^. Exploiting this efficiency, PlantCAD2 takes 8,192 base pair (bp) windows, which is a 16-fold increase over the 512-bp windows used in PlantCAD. Second, to evaluate the effect of model sizes on performance, we trained a series of depth-scaled PlantCAD2 models of 88M, 311M and 694M parameters (**Figure 1A-1B**), which we named PlantCAD2-S, PlantCAD2-M, and PlantCAD2-L respectively. As expected, following pre-training, the largest model (PlantCAD2-L) demonstrated the best masked token prediction accuracy (0.657), followed by PlantCAD2-M (0.641) and PlantCAD2-S (0.598), when evaluated on hold-out test set by randomly masking 15% of nucleotides per sequence (**Figure S1B**). However, the largest model also shows slowest inference speed at ∼32,238 nucleotides per second on a NVIDIA H100 80 GB PCIe GPU (**Figure 1C**) and uses ∼27G GPU memory with a of 32 (**Table 3**), reflecting the typical trade-off between accuracy and computational efficiency. The smallest PlantCAD2 model, PlantCAD2-S, achieves a 4.5-fold (**Figure 1C**) higher inference throughput while requiring considerably less GPU memory at equivalent batch sizes (**Table 3**), highlighting its practicality for high-throughput analyses. Despite differences in model size, the three models showed high correlation in their per-species prediction accuracies (r > 0.97), suggesting consistent learning patterns across scales (**Figure S1C; Supplemental Table 1**). Third, to assess the effect of input length on pretraining accuracy, we varied the context window size from 512bp to 8,192bp and evaluated performance by masking the central token. All three models showed improved masked token prediction accuracy with longer contexts, underscoring the importance of extended context for modeling kilobase-scale genomic dependencies (**Figure 1D**). Lastly, we expanded the evolutionary diversity of the training dataset from 16 to 65 angiosperm genomes (**Figure 1E**; **Supplemental Table 1**), selecting one representative species per genus to maximize phylogenetic breadth. When analyzing pre-training performance across species, we found a weak positive correlation between genome size and masked token accuracy (r = 0.525; **Figure S1D, Supplemental Table 1**). This relationship is likely driven by the fact that larger genomes tend to contain more repetitive sequences ^46^. Since the masked language modeling objective can predict repetitive elements more easily than non-repetitive elements even after applying down-sampling and down-weighting (see Methods), the amount of repeats in the test set could inflate accuracy ^22,27,35^. Consistent with our expectation, we also detected a positive correlation between the number of repeats in the test set and masked language modeling accuracy (**Figure S1E**), this also highlights the importance of down-sampling and down-weighting repetitive sequences ^35^ in pre-training DNA language models.

**Figure 1.**
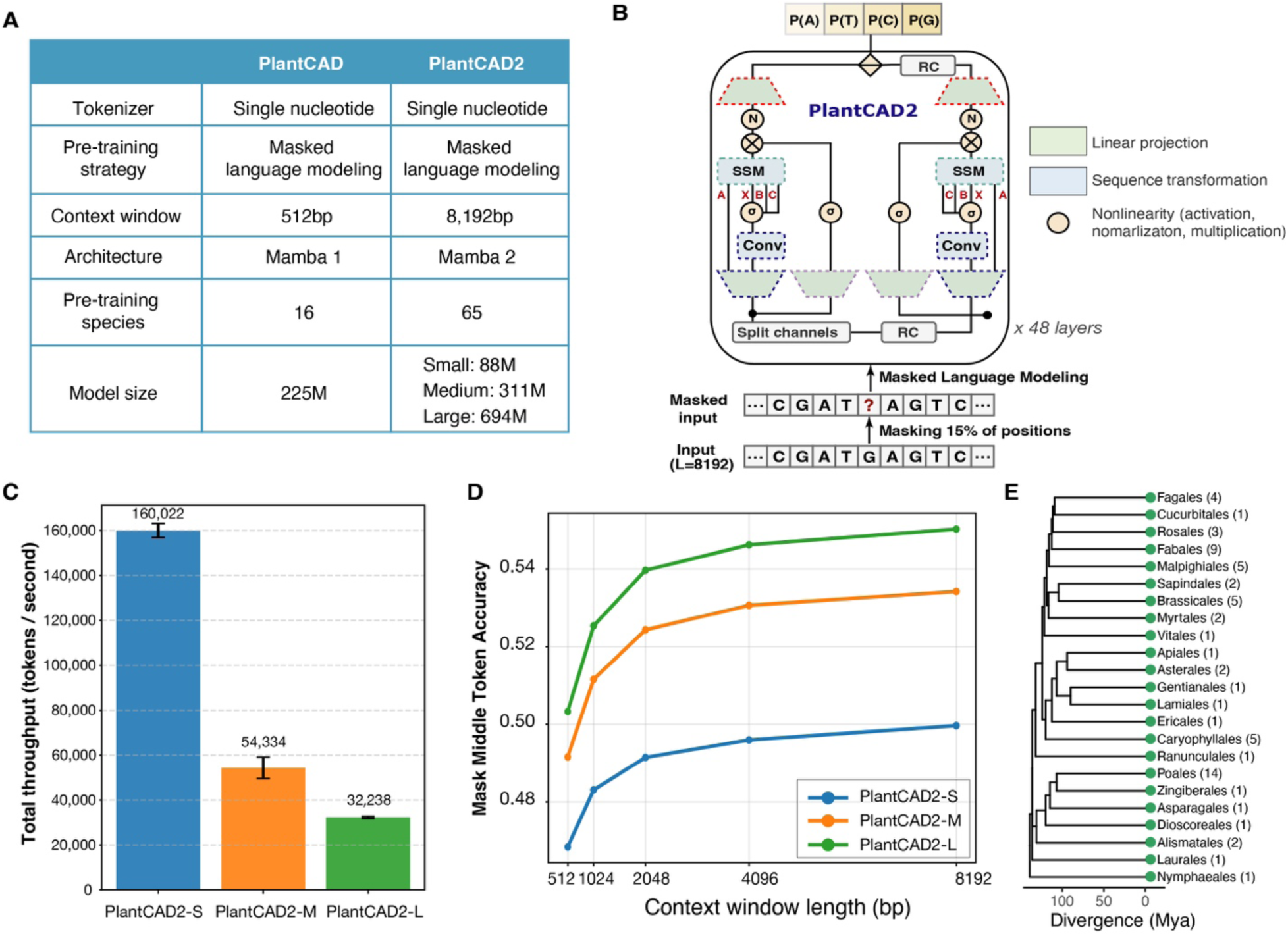
Overview of the PlantCAD2 model. **(A)** Comparison of PlantCAD1 and PlantCAD2 model configurations. PlantCAD2 introduces a longer context window, upgraded architecture (Mamba2), expanded pre-training species set, and scaled model sizes (small: 88M, medium: 311M, large: 694M parameters), while maintaining single-nucleotide tokenization. **(B)** Schematic of the PlantCAD2 architecture based on Mamba2 with reverse-complement (RC) equivariance, convolutional and state space modules (SSM), and a masked language modeling objective applied to 8,192 bp input sequences. **(C)** Total throughput (sequences/second) of PlantCAD2 models on NVIDIA H100 80 GB PCIe GPU across batch sizes (1–64). Values on the bar represent mean throughput across batch sizes. **(D)** Effect of context window length on model performance. The y-axis shows the prediction accuracy of three models when masking the single central token in the held-out test set. **(E)** Phylogenetic distribution of the 65 angiosperm genomes across flowering plant orders. Numbers in parentheses indicate the number of species included from each order.

**Table 3.** Inference throughput and GPU memory usage of PlantCAD2 models on an NVIDIA H100 80 GB PCIe GPU.

| Model | Seq Len<br>(bp) | Batch size | Peak<br>memory<br>(GB) | Seq/s | Tokens/s |
| --- | --- | --- | --- | --- | --- |
| PlantCAD2-S | 8192 | 1 | 0.79 | 19.58 | 160,422 |
|  |  | 4 | 1.94 | 20.12 | 164,812 |
|  |  | 8 | 3.55 | 19.63 | 160,829 |
|  |  | 16 | 6.56 | 19.61 | 160,653 |
|  |  | 32 | 12.76 | 19.26 | 157,767 |
|  |  | 64 | 24.89 | 19 | 155,649 |
| PlantCAD2-M | 8192 | 1 | 2.5 | 5.46 | 44,761 |
|  |  | 4 | 3.49 | 7.02 | 57,489 |
|  |  | 8 | 5.54 | 6.88 | 56,388 |
|  |  | 16 | 9.62 | 6.88 | 56,386 |
|  |  | 32 | 17.62 | 6.76 | 55,342 |
|  |  | 64 | 33.62 | 6.79 | 55,636 |
| PlantCAD2-L | 8192 | 1 | 4.76 | 3.98 | 32,634 |
|  |  | 4 | 5.75 | 4.01 | 32,847 |
|  |  | 8 | 8.83 | 3.94 | 32,264 |
|  |  | 16 | 14.89 | 3.92 | 32,111 |
|  |  | 32 | 26.95 | 3.87 | 31,741 |
|  |  | 64 | 51.09 | 3.89 | 31,833 |

### PlantCAD2 accurately predicts evolutionary conservation with a zero-shot strategy

Evolutionary conservation, commonly estimated from multiple sequence alignment (MSA), is widely used to identify deleterious mutations that may reduce organismal fitness ^47–50^. However, estimating a genome-wide MSA is particularly challenging in plants due to extensive transposable element (TE) insertions and their high turnover rate, which obscure orthologous relationships outside of conserved coding regions ^51^. This limitation highlights the need for alignment-free approaches to assess conservation across diverse plant genomes. Given that PlantCAD2 is pre-trained on 65 evolutionary distant species, we hypothesize that PlantCAD2 can be used to predict evolutionary conservation without an MSA. We first evaluated how accurate PlantCAD2 is to distinguish highly conserved sites versus less conserved sites using a zero-shot strategy. As illustrated in **Figure 2A**, we used the masked nucleotide/token prediction accuracy from the frozen model to represent per-base conservation, which means highly conserved bases would receive higher predicted probabilities for the reference allele, whereas less conserved bases would yield lower confidence scores. We benchmarked the performance of PlantCAD2 against three baselines: its predecessor PlantCAD ^27^, GPN ^22^ (a plant specific DNA LM trained on Brassicales genomes), and Evo2 ^19^, a general-purpose DNA language model pre-trained using a causal language modeling (CLM) objective, also known as next-token prediction. Unlike masked language modeling, which enables access to both upstream and downstream context, CLM imposes a strict left-to-right constraint, meaning Evo2 is inherently unidirectional. Therefore, we input the entire sequences without masking for Evo2 and use the likelihood of the model to represent conservation. Notably, Evo2 was trained at a substantially greater scale, with 7 billion parameters and 9.3 trillion nucleotides, which is over 310 times more training data than used for PlantCAD2, therefore providing a rigorous benchmark for assessing the efficiency and representational power of our models. We excluded AgroNT from zero-shot evaluation as its non-overlapping k-mer tokenization strategy prevents single-nucleotide resolution tasks, and we previously demonstrated its limited zero-shot capabilities ^27^.

**Figure 2.**
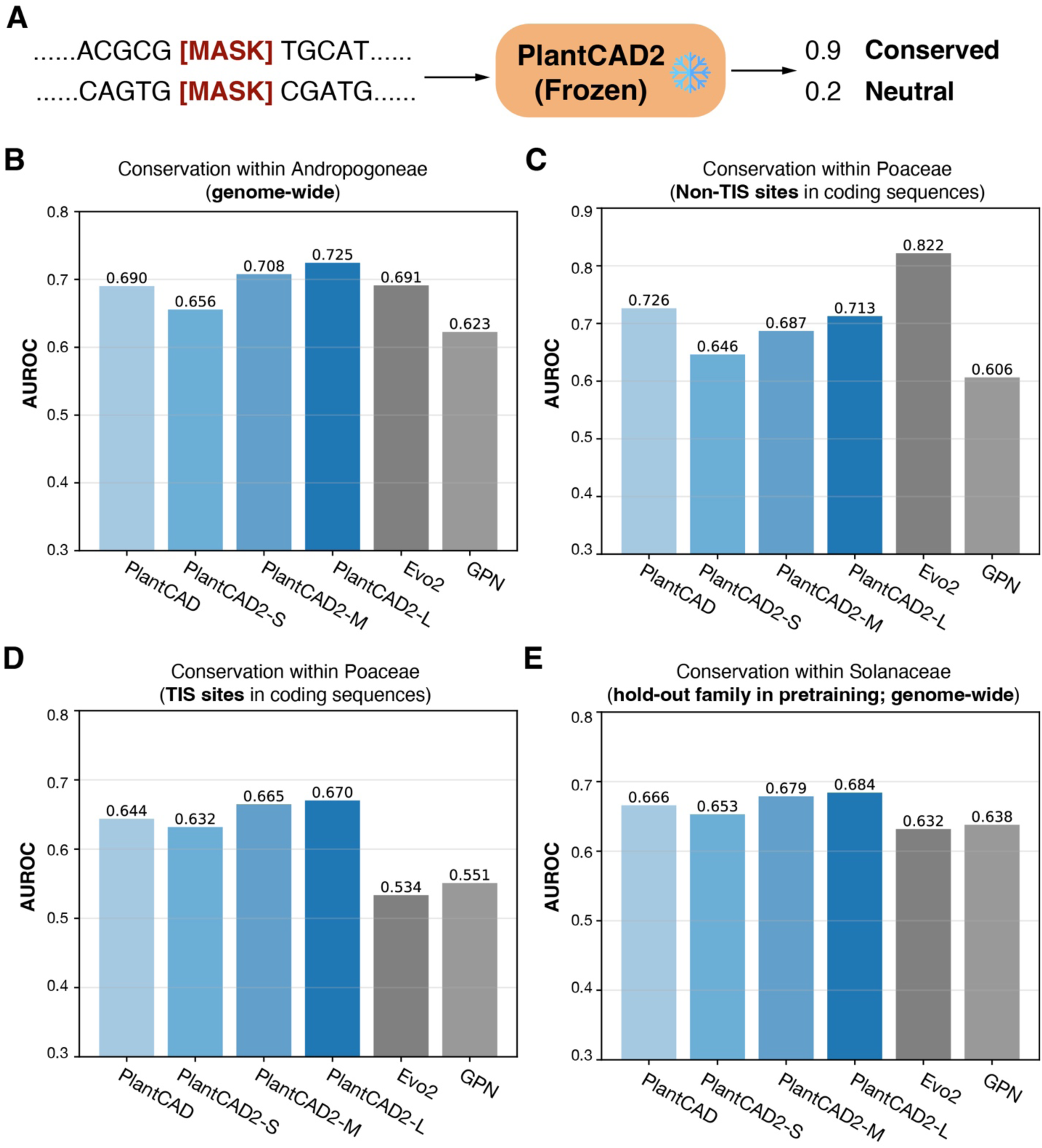
PlantCAD2 accurately predicts evolutionary conservation using zero-shot strategy. **(A)** Illustration of the zero-shot conservation approach. For each masked token (nucleotide) in the reference sequence, the frozen PlantCAD2 model outputs the probability of the reference allele; higher probabilities indicate higher evolutionary constraint. **(B)** AUROC for predicting conserved versus neutral sites in sorghum, representing conservation within the Andropogoneae tribe. **(C-D)** AUROC for predicting conservation within Poaceae using sites in coding regions, evaluated separately for non-TIS **(C)** and TIS **(D)**. **(E)** AUROC for conservation prediction within Solanaceae using SNPs from potato. Solanaceae species were entirely excluded from the PlantCAD2 pre-training, making this a hold-out family used to evaluate cross-lineage generalization.

We assessed this strategy in three independent tasks. First, we performed cross-species alignments of 34 Andropogoneae genomes ^27^ to the sorghum reference genome, and identified highly conserved and less conserved sites based on alignment coverage and identity (see Methods). PlantCAD2 consistently outperformed PlantCAD in distinguishing highly conserved from less conserved sites in the sorghum genome, with the largest PlantCAD2 achieving the highest AUROC (**Figure 2B**; **Supplemental Table 2**). Notably, PlantCAD2-M achieved slightly better performance than Evo2 (AUROC 0.708 vs 0.691) despite being ∼22-fold smaller (311M vs 7B parameters), while PlantCAD2-L, being ∼11-fold smaller (694M parameters), further improved to 0.73. This demonstrates that our PlantCAD2 models can match or exceed Evo2’s performance with far fewer parameters. Given that PlantCAD2 is pre-trained with a context window of 8192 bp, we also examined the effect of context length on conservation prediction. AUROC scores increased with longer sequence contexts, plateauing at 4096 bp for all PlantCAD2 models (**Figure S2A; Supplemental Table 2**). These findings indicate that evolutionary constraint signals benefit from broader sequence context and that larger models with extended receptive fields are better suited to capture these dependencies.

In the second task, we used multiple sequence alignments from coding sequences of 325 Poaceae genomes to calculate phyloP scores and define highly conserved sites (phyloP > 5) and less conserved sites (phyloP < 1.5). While the relationship between selection and phyloP scores can be nuanced ^52^, restricting phyloP calculation to coding regions helps mitigate alignment noise caused by the very high transposable element turnover rate in plant genomes ^53^, providing a more reliable benchmark for conservation prediction. Given Evo2’s unidirectional nature, we hypothesized it might struggle with features requiring bidirectional context, particularly translation initiation sites (TIS), where both upstream regulatory motifs in the 5′ UTR and downstream coding sequence context critically influence start codon recognition and conservation ^54^. To test this hypothesis, we separately evaluated performance on TIS versus non-TIS positions within coding sequences. For non-TIS sites, PlantCAD2 models showed lower performance compared to Evo2 (**Figure 2C**; **Supplemental Table 2**), potentially because Evo2’s training data included mature mRNA sequences while PlantCAD2 was trained exclusively on genomic DNA, giving Evo2 an advantage in coding sequence conservation tasks. We also note that the original PlantCAD slightly outperforms all PlantCAD2 models on this specific task (**Figure 2C**). This likely reflects differences in input–pretraining alignment: PlantCAD was pre-trained on short 512 bp windows enriched for coding regions, and the CDS-centered inputs used here closely match that training setting. In contrast, PlantCAD2 was pre-trained on 8 kb genomic windows that systematically include intronic and intergenic contexts. Consequently, restricting the input to CDS-centered sequences removes intronic context and deviates from PlantCAD2’s pre-training setting, attenuating the advantages conferred by its long receptive field. Nevertheless, despite this mismatch, PlantCAD2 models still achieve competitive performance, indicating that their learned representations remain informative even under constrained input conditions. Interestingly, when evaluating TIS conservation, we observed a strong bias in Evo2: its AUROC dropped drastically to 0.534, barely above random. In contrast, PlantCAD2 maintained robust performance (AUROC: 0.632–0.670; **Figure 2D**; **Supplemental Table 2**). Even the 65M-parameter GPN outperformed the 7B-parameter Evo2 on this task (AUROC of 0.551), further highlighting Evo2’s architectural limitations for TIS prediction (**Figure 2D**; **Supplemental Table 2**). This TIS-specific weakness in Evo2 validates our hypothesis: without access to coding sequence contexts that are more evolutionary constrained, Evo2 cannot properly assess the conservation patterns at translation start sites. Consistent with the previous task, we observed only modest improvements from longer context windows, indicating that evolutionary conservation signals are predominantly local (**Figure S2B-S2C**). These results demonstrate that PlantCAD2 provides more consistent and unbiased conservation predictions across different genomic contexts.

To further assess whether PlantCAD2 generalizes to lineages absent from pre-training. We evaluated its zero-shot conservation predictions in Solanaceae, a major angiosperm family not represented in the 65 pre-training species. This evaluation used SNPs from the potato (*Solanum tuberosum* L.) genome, with GERP scores ^55^ calculated from a multiple sequence alignment of 95 Solanaceae genomes ^56^. Conserved sites were defined as SNPs with GERP scores greater than 1.5, and neutral sites as SNPs with GERP scores lower than 0. As shown in **Figure 2E**, PlantCAD2 models achieved higher AUROC than both Evo2 and GPN, despite multiple Solanum genomes being part of the Evo2 pre-training corpus but absent from PlantCAD2 pre-training. Additionally, to understand functional constraints across the entire genome, we stratified the evaluation into six distinct genomic contexts: CDS, upstream (promoters), downstream, introns, intergenic regions, and transposable elements (TEs) (**Figure S3**). As expected, accuracy was higher in coding versus non-coding regions (**Figure S3**), likely because the stronger functional constraints on CDS make these patterns easier for the model to capture. When comparing models across the five non-coding categories, PlantCAD2 consistently showed the best performance in four categories (upstream, downstream, intergenic, and TEs), with GPN showing only a marginal advantage in introns. Notably, PlantCAD2 consistently surpassed Evo2 across all six genomic contexts. The strong consistent performance across three phylogenetically distant angiosperm lineages indicates that PlantCAD2 captures fundamental conservation principles shared across flowering plants rather than lineage-specific patterns. This enables accurate alignment-free conservation inference even in lineages unseen during pre-training.

### PlantCAD2 accurately predicts within-species conserved transcriptional and translational junction sites with zero-shot strategy

We next quantified how well PlantCAD2 captured the sequence context that defines key transcriptional and translational junctions. Using a similar zero-shot strategy, we designed four tasks to recapitulate motifs (**Figure 3**). Instead of masking one base pair, for each annotated junction, we replaced the annotated motif with consecutive [MASK] tokens. Given the non-canonical motifs do exist in plants ^57^, we retained all annotated sites without filtering for canonical signals. Consequently, the evaluation included both canonical and non-canonical variants as defined in the reference annotation. For example, translation initiation sites included both canonical ATG and non-canonical start codons such as GTG and TTG. We then extracted a fixed 8,192-bp window centered on the masked motif and presented the entire masked sequence to the model without fine-tuning. A prediction was considered correct if the model’s top-1 reconstruction exactly matched the ground truth motif found in the genome.

**Figure 3.**
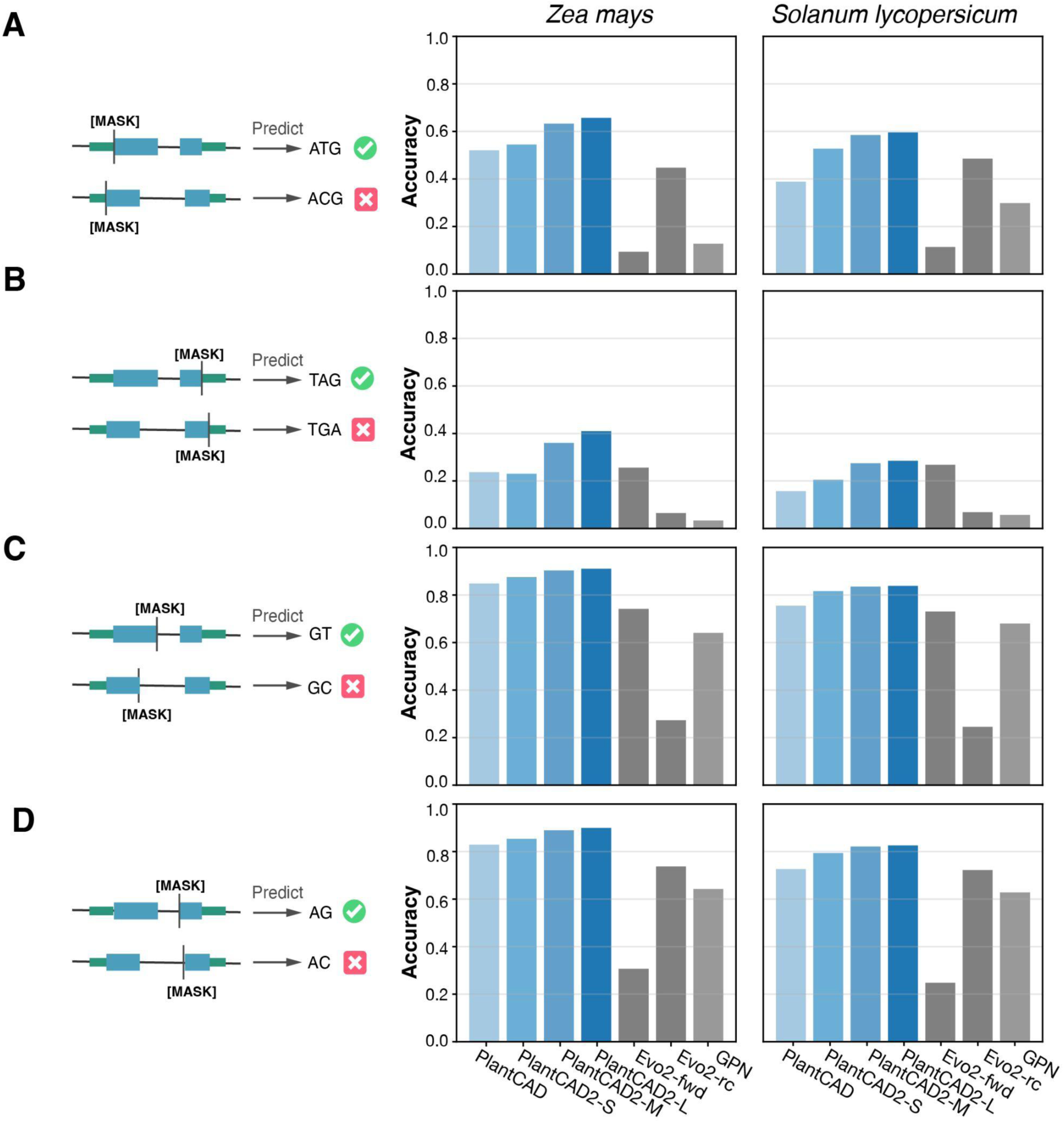
PlantCAD2 accurately predicts transcriptional and translational junction sites using zero-shot masked motif prediction. Left panels show the masking strategy where canonical motifs are replaced with [MASK] tokens and models predict the correct sequence. Right panels show prediction accuracy for each model on maize (left, included in training) and tomato (right, excluded from training). **(A)** Translation initiation sites (ATG masking). **(B)** Translation termination sites (TAG/TGA/TAA masking). **(C)** Splice donor sites (GT masking). **(D)** Splice acceptor sites (AG masking).

As above, we benchmarked the performance of PlantCAD2 against PlantCAD, GPN and Evo2. Given what we observed in **Figure 2D** that Evo2 is limited with its poor TIS conservation prediction, we evaluated it using two configurations: (1) forward sequences (Evo2-fwd), where the model uses upstream context to predict the junction, and (2) reverse-complement sequences (Evo2-rc), where the model uses downstream context (reverse complemented) to predict the junction. For GPN and PlantCAD, which are both limited to context windows of 512 bp, we used 512-bp windows centered on the junctions for evaluation.

When evaluated on both maize (*Zea mays*), which was included in pre-training and tomato (*Solanum lycopersicum*), which was not included in pre-training, PlantCAD2 consistently outperformed PlantCAD1 across all junction types (**Figure 3**; **Supplemental Table 3**). Notably, even the smallest PlantCAD2 model (88M parameters) outperformed the original PlantCAD (311M parameters), demonstrating that architectural improvements and expanded phylogenetic diversity (65 vs. 16 genomes) provide substantial benefits beyond parameter scaling alone. Accuracy increased with model size, following expected scaling law, with the largest model achieving the highest masked-motif prediction accuracy across both species.

As expected, Evo2 showed strong directional effects. For junctions where downstream context is more informative (TIS and splice acceptor), Evo2-rc performed better, as the reverse-complement orientation allows the model to ’see’ the downstream coding sequences that provide stronger signals. Conversely, for junctions where upstream context matters more (TTS and splice donor), Evo2-fwd showed superior performance. The sharp contrast in performance between forward sequences and reverse complemented sequences reflects a fundamental limitation of causal language models: their unidirectional nature prevents them from accessing both upstream and downstream signals simultaneously. In contrast, PlantCAD2’s bidirectional and reverse-complement equivariant design achieved robust performance regardless of sequence orientation, consistently leveraging both upstream and downstream context for all junction types.

While recovering canonical junction motifs demonstrates basic sequence understanding, we next tested whether PlantCAD2 captures deeper evolutionary signals that distinguish core genes (evolutionarily constrained and present across taxa) from non-core genes (rapidly evolving and taxa-specific). We extracted each model’s log-likelihood of the canonical motif as a conservation score and evaluated binary classification performance using AUROC (**Figure S4**). For Evo2, we selected the optimal orientation based on junction type (forward for TTS/donor, reverse-complement for TIS/acceptor) as determined above. Remarkably, even though the tomato genome was excluded from PlantCAD2’s 65 pre-training genomes, PlantCAD2 consistently outperformed Evo2—which did include tomato during pre-training. This demonstrates strong cross-species generalization: PlantCAD2 learned transferable conservation patterns from other angiosperms that effectively predict functional constraints in unseen species. These results highlight PlantCAD2’s ability to capture fundamental evolutionary principles instead of just recognizing simple motif recognition.

### PlantCAD2 predicts functional structural variants with zero-shot strategy

In addition to single-nucleotide conservation, we investigated whether PlantCAD2 can generalize to predicting the functional impact of structural variants, such as small deletions, using a zero-shot approach. To do this, we simulated a set of deletions in the Arabidopsis genome and computed the Δlog P score, defined as the log-likelihood ratio between the mutated and reference sequences surrounding the deletion site, averaged across the deletion window (**Figure 4A**). To assess how well Δlog P reflects the functional deletions, we utilized two complementary conservation metrics derived from multiple sequence alignments of 63 genomes ^58^: phyloP ^59^, which measures conservation or acceleration at individual sites, and phastCons ^59^ which identifies conserved elements by analyzing patterns across adjacent sites. We classified deletions as either "highly conserved" or "less conserved" based on the average scores from each metric.

**Figure 4.**
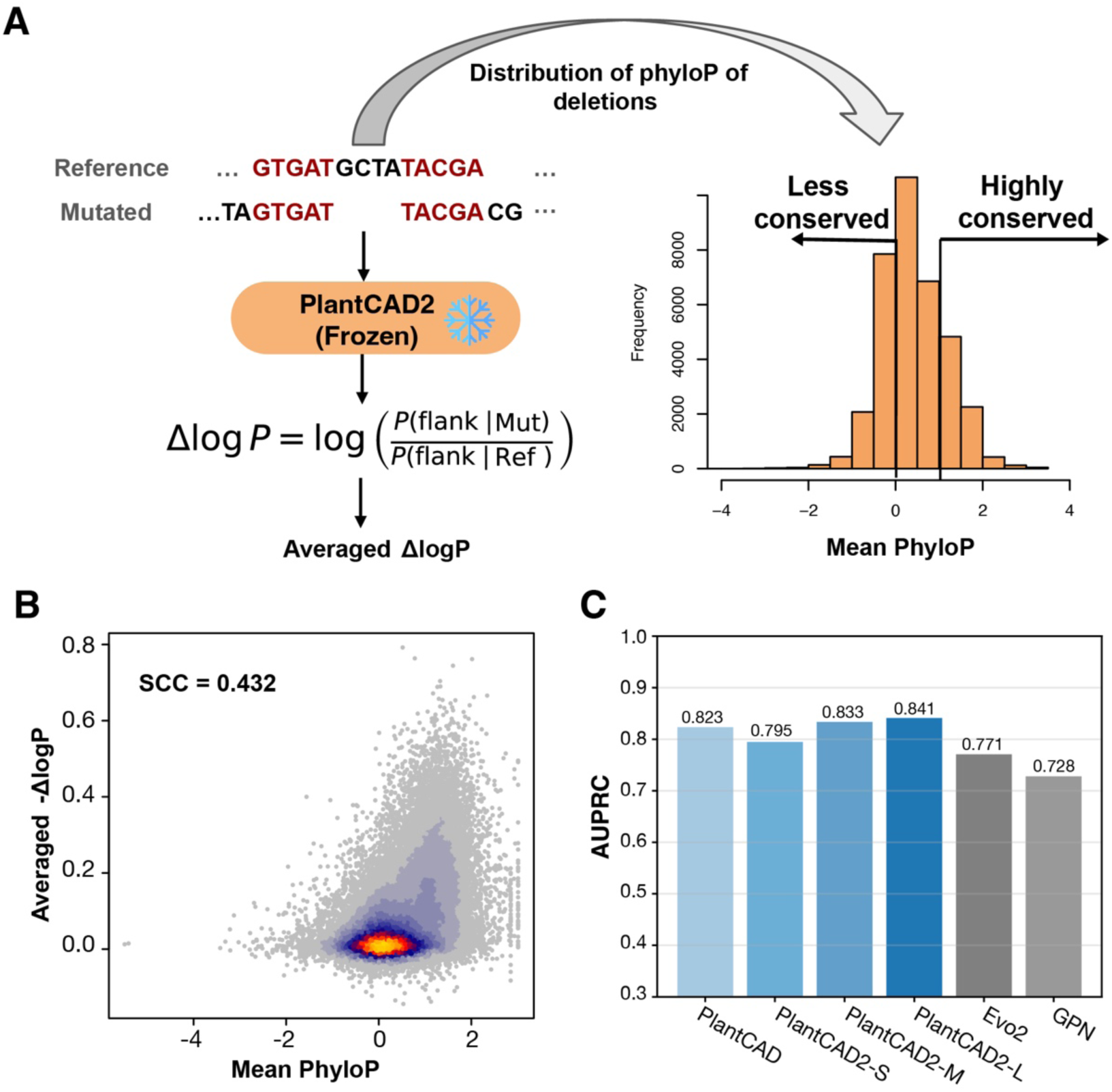
PlantCAD2 predicts functional impact of structural variants using zero-shot strategy. **(A)** ΔlogP calculation approach for deletion variants and phyloP score distribution for classification. **(B)** Scatter plot showing the positive correlation between PlantCAD2’s -ΔlogP scores and phyloP-based conservation scores. **(C)** AUROC performance distinguishes highly conserved from less conserved deletions.

Given more negative Δlog P represents higher conservation, we observed PlantCAD2’s negative Δlog P scores showed strong positive correlation with both phyloP-based and phastCons constraint scores (**Figure 4B**; **Figure S5**), with the model assigning higher likelihoods to mutations in evolutionarily constrained regions. To quantify this relationship, we binarized deletions into "highly conserved" and "less conserved" categories based on either phyloP (**Figure 4A**) or phastCons scores (**Figure S5A**) and evaluated whether Δlog P could discriminate between them. PlantCAD2 achieved robust classification performance either using phyloP (**Figure 4C**; **Supplemental Table 4**) or phastCons (**Figure S5**), with even the 88M-parameter PlantCAD2-S outperforming the 7B-parameter Evo2, highlighting the advantage of plant-specific training over general-purpose models again. Additionally, we also observed classification performance is saturated with just 20 bp of flanking sequence on each side (**Figure S6**), indicating that local sequence context sufficiently captures the functional impact of small deletions. This strong performance is particularly impressive given that PlantCAD2 was never explicitly trained on structural variants, suggesting it learned generalizable sequence constraint patterns during pre-training. These results suggest that PlantCAD2’s learned representations generalize beyond single-nucleotide changes, capturing broader sequence dependencies relevant to noncoding structural variation. This provides a scalable alternative to traditional alignment-based conservation methods. Notably, this also represents one of the first efforts to use DNA LMs for estimating indel effects in plant genomes, underscoring the potential of foundation models in addressing complex variant interpretation challenges.

### Transcription factor binding sites revealed by high-confidence predictions of PlantCAD2

To evaluate the interpretability and regulatory grammar capture of PlantCAD2, we first assessed whether the model’s high-confidence predictions align with experimental verified transcription factor binding sites (TFBS) using a zero-shot approach ^22^. We masked each position within DAP-Seq-defined DREB2 peak regions ^60^ and generated importance scores from the model’s output distribution, then clustered these into motifs using TF-MoDISco ^61^. DREB2 (AP2/ERF family) was selected as a representative transcription factor because AP2/ERF TF family is prevalent and evolutionary conserved across angiosperms ^62^.

The de novo motif discovered by TF-MoDISco with PlantCAD2 predictions showed a highly significant match to the canonical AP2/ERF profile (https://jaspar.elixir.no/matrix/MA1222.1) in the JASPAR database (Tomtom p-value = 0.0001; **Figure S7**). In contrast, while the Evo2 model also detected the motif, the match was substantially less significant (Tomtom p-value=0.005), suggesting that PlantCAD2 captures plant-specific regulatory features with higher fidelity.

We then extended this strategy to performing de novo motif discovery across all 1.5-kb regions upstream of translation initiation sites in the Arabidopsis genome. This genome-wide analysis was designed to systematically identify representative cis-regulatory motifs captured by the pre-trained PlantCAD2 model beyond known DAP-Seq peaks. In total, 104 high-confidence motifs were identified. Of the 104 high-confidence motifs, 27 matched known entries in the JASPAR plant database (Tomtom p < 0.05). This 26% match rate likely underestimates the true proportion of functional motifs, given that JASPAR’s plant subset covers only a fraction of known plant transcription factor families and is heavily biased toward a few model organisms. Representative examples, highlighted in **Figure S8** include BPC1 (AT2G01930), involved in ovule identity ^63^; TSO1 (AT3G22780), a regulator of floral cell division ^64^; SRS7/SPRI2 (AT1G19790), which modulates auxin-dependent reproductive barriers ^65^; and ZML1 (AT3G21175), a GAGA-type factor mediating photoprotection ^66^. These results may suggest these TFBS exhibit profound evolutionary conservation and indicate PlantCAD2 could be a useful tool to recover both established and potentially novel conserved regulatory elements. All discovered motifs and their statistics are available on GitHub: https://plantcad.github.io/TFBS.

### Fine-tuning PlantCAD2 accurately predicts cross-species chromatin accessible regions and cell-type-specific accessible regions

We next investigated whether PlantCAD2 learned chromatin states by assessing its performance in predicting genome-wide chromatin accessibility across multiple plant species. We formulated this as a binary classification task, in which the model predicts whether a given genomic region corresponds to an accessible chromatin region, as defined by ATAC-seq (**Methods**). We used recently published ATAC-seq data including 11 diverse plant species, including both dicots and monocots ^67^. In this task, positive examples correspond to accessible peaks from ATAC-Seq, while negative examples were from genomic background regions. We used 600-bp genomic windows for all models, as this resolution captures the typical size of ATAC-seq peaks while providing sufficient sequence context for regulatory element prediction. Due to the biological reality of accessible regions comprising only a small fraction of the entire genome, this task is highly imbalanced (**Supplementary Table 5**). For example, less than 1% of regions in the maize genome are labeled as positive.

To effectively leverage the pre-trained foundation model, we used a Low-Rank Adaptation (LoRA) ^68^ fine-tuning strategy for PlantCAD2, which inserts small trainable rank-decomposition matrices into the feedforward layers while keeping the rest of the model frozen (**Figure 5A**). This approach updates only a small fraction of parameters, enabling efficient task-specific adaptation with minimal risk of overfitting or forgetting the pre-trained knowledge. To assess the contribution of pre-training, we compared this approach to two supervised baselines: (1) a fully supervised version of PlantCAD2-S from scratch, where all model parameters were randomly initialized and updated during training; and (2) a commonly used CNN+LSTM [33] architecture trained from scratch. We also benchmarked against AgroNT ^24^, another plant-specific DNA LM. We excluded GPN and PlantCAD due to their limited 512-bp context window. For other DNA LMs, we tailored our benchmarking strategy to the specific characteristics of each model. Specifically, we benchmarked against AgroNT using LoRA, as our previous study ^27^ demonstrated that AgroNT yields suboptimal performance with frozen embeddings but gains significant accuracy through fine-tuning, which is computationally feasible given its 1-billion-parameter size. In contrast, while fine-tuning the 7-billion-parameter Evo2 model was computationally expensive, we sought to strictly evaluate the quality of its learned representations against PlantCAD2. Therefore, we implemented a dual-probing strategy using frozen embeddings extracted from the last hidden state of both models. To ensure a rigorous comparison, we evaluated embeddings derived from three distinct input configurations: the forward sequence, the reverse-complement sequence, and the average of both. This comprehensive approach ensures that we captured the maximal information content available from Evo2, accounting for potential strand biases or invariant features. We then trained two distinct classifiers: (1) a simple linear layer to assess linear separability; and (2) XGBoost ^69^ to capture complex non-linear feature interactions. This ensures a comprehensive evaluation of the information content encoded by each model without requiring full parameter updates. All models were trained using Arabidopsis and validated on hold-out chromosomes within Arabidopsis as well as on 10 additional test species spanning a broad phylogenetic range.

**Figure 5.**
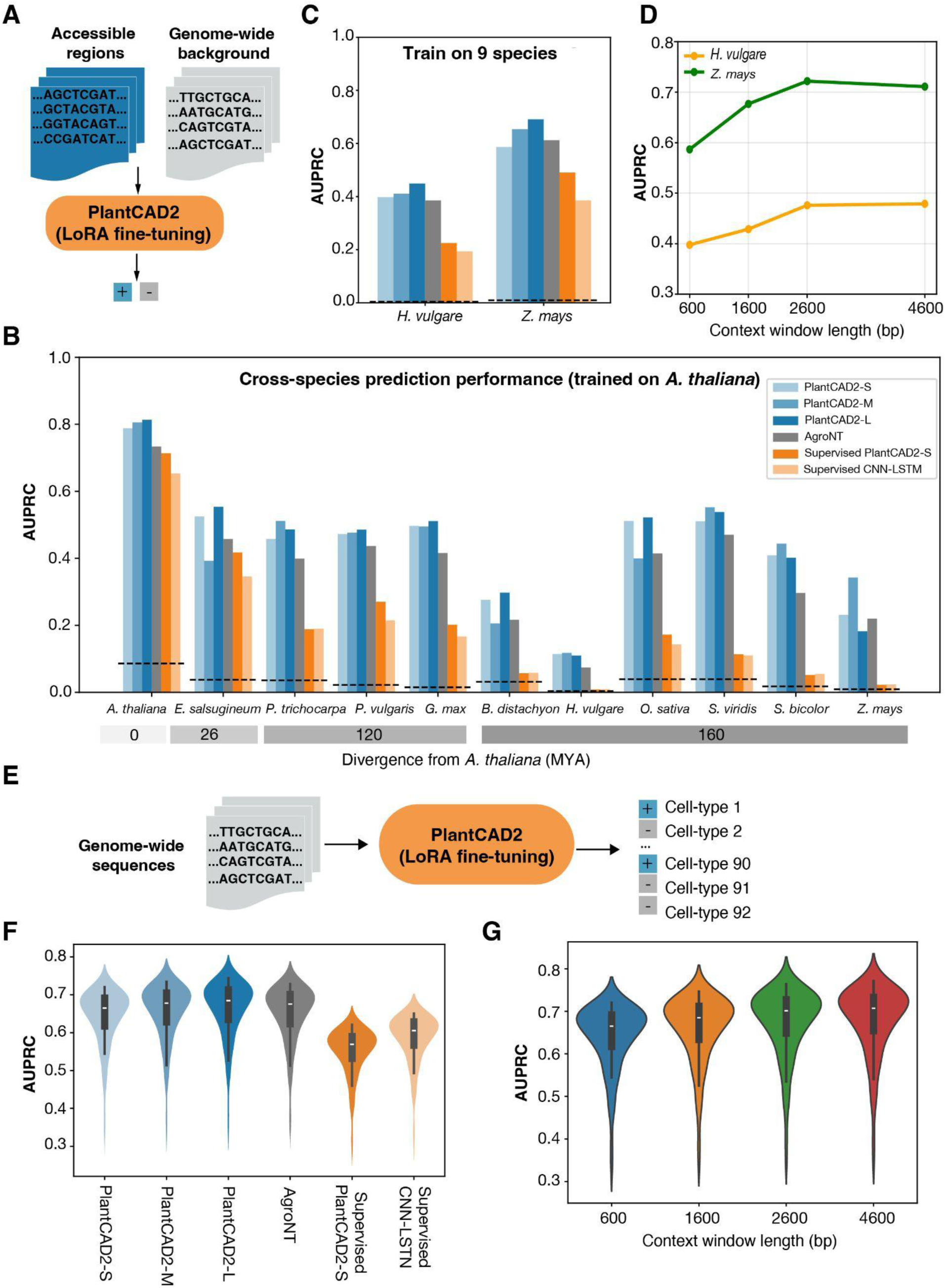
PlantCAD2 predicts chromatin accessibility across species and cell types. **(A)** LoRA fine-tuning approach for binary accessibility prediction using ATAC-seq peaks versus genomic background. **(B)** Cross-species AUPRC performance when trained on Arabidopsis, showing superior generalization of PlantCAD2 models compared to supervised baselines across evolutionary distances. **(C)** Multi-species training performance on held-out barley and maize. **(D)** Effect of context window length on accessibility prediction accuracy for PlantCAD2-S. **(E)** Multi-label classification approach for cell-type-specific accessibility prediction. **(F)** Performance comparison across models for 92 cell types . **(G)** Context window effects on cell-type-specific prediction accuracy for PlantCAD2-S.

Given the strong class imbalance of this task, we measured model performance using the area under the precision–recall curve (AUPRC), which is more informative than AUROC in imbalanced classification settings. LoRA fine-tuned PlantCAD2 consistently achieved the best performance in both within-species evaluation and cross-species generalization, outperforming supervised baselines and AgroNT across all test species (**Figure 5B**; **Supplementary Table 5**). Furthermore, in the comparison using fixed (frozen) embeddings against the 7-billion-parameter Evo2, PlantCAD2 consistently achieved higher AUPRC scores across the 11 species, regardless of whether a linear or non-linear classifier was used (**Figure S9**). While the averaged embedding (combining forward and reverse-complement strands) proved to be the most robust configuration for Evo2, it was still consistently outperformed by all PlantCAD2 models across all test species. Remarkably, even our smallest model, PlantCAD2-S, surpassed the 7B-parameter Evo2. On average, PlantCAD2-L yielded an AUPRC improvement of 0.1142 and 0.1735 over Evo2’s best performance using Linear and XGBoost classifiers, respectively (**Figure S9**). This result highlights the efficiency of PlantCAD2 as an angiosperm DNA foundation model, demonstrating that domain-specific pre-training contributes more to representation quality than simply scaling up model parameters.

Across the diverse species tested, we observed a strong negative correlation between genome size of AUPRC, which reflects the increasing difficulty of distinguishing sparse regulatory elements in large intergenic regions (**Figure S10**). Despite this challenge, fine-tuned DNA LMs consistently outperformed supervised models trained from scratch (including CNN+LSTM and Supervised PlantCAD2-S), indicating that pre-training enables better learning of chromatin states, particularly when transferring knowledge across species (**Figure 5B**). Specifically, the supervised models (whether using CNN+LSTM or Supervised PlantCAD2-S) trained on Arabidopsis generalized reasonably well to closely related dicots, but their performance declined substantially when applied to evolutionarily distant monocots such as maize and barley. In contrast, fine-tuned PlantCAD2 retained strong predictive accuracy across both dicots and monocots, demonstrating its ability to capture regulatory features conserved across deep evolutionary divergence. These results underscore the power of combining self-supervised pre-training with parameter-efficient fine-tuning for plant regulatory genomics.

While the foundation model fine-tuned on Arabidopsis already demonstrated clear advantages in cross-species prediction, we further fine-tuned a multi-species version of PlantCAD2 using accessible chromatin regions from multiple plant genomes to enhance its robustness across diverse lineages. With maize and barley held out as test species, this multi-species model achieved impressive AUPRC scores of 0.691 for maize and 0.449 for barley (**Figure 5C**; **Supplemental Table 6**). To investigate whether extended sequence context could further improve prediction accuracy, we maintained the same 600-bp labels but varied the input window size by including different amounts of flanking sequence. For computational efficiency, we conducted this analysis using PlantCAD2-S. Performance consistently improved with longer context windows, with AUPRC increasing from 0.587 to 0.711 for maize and from 0.398 to 0.479 for barley when extending from 600 bp to 4,600 bp (**Figure 5D**). This substantial improvement suggests that distal regulatory elements and broader chromatin context beyond the immediate peak boundaries contribute to accessibility prediction, highlighting the advantage of PlantCAD2’s long-context architecture. These fine-tuned models serve as a robust predictor of chromatin accessibility across flowering plants and are publicly available to the community as a ready-to-use resource for regulatory annotation in non-model species.

To further assess whether PlantCAD2 can resolve cell-type–specific regulatory landscapes, we tested its ability to predict accessible chromatin regions identified through single-cell ATAC-seq (scATAC-seq) in maize ^70^. In contrast to the binary classification task used for genome-wide accessibility, this task was framed as a multi-label classification problem, where each genomic window could be accessible in one or more cell types (**Figure 5E**). We curated high-confidence cell-type–specific peaks across major maize tissues from published scATAC-seq datasets (**Methods**), using them as labels for multi-label fine-tuning and evaluation.

We applied the same LoRA fine-tuning strategy used in prior experiments, adapting PlantCAD2 to predict cell-type–specific accessibility using only a small number of trainable parameters. As in previous sections, we compared performance against two supervised baselines: a CNN+LSTM model trained from scratch and a fully supervised version of PlantCAD2. All models were trained on all maize cell types and evaluated on held-out chromosomes, with performance measured using micro-averaged precision-recall curves across cell types. Despite the complexity and subtlety of cell-type–specific regulatory signatures, LoRA fine-tuned PlantCAD2 can still achieve very high accuracy and outperformed other baselines (**Figure 5F**; **Supplemental Table 7**). Similar to our genome-wide accessibility results, extending the input context window beyond the core 600-bp peak region further improved cell-type specificity, with AUPRC increasing from 0.665 to 0.707 when using 4,600-bp windows (**Figure 5G**). This suggests that cell-type–specific regulatory programs are influenced by broader chromatin context and distal regulatory interactions. The model captured both shared and lineage-specific accessibility patterns, demonstrating that pre-trained DNA representations can be effectively adapted to fine-grained regulatory annotations. These results suggest that PlantCAD2 is not only effective at modeling general chromatin accessibility across species but is also capable of distinguishing cell-type–specific regulatory programs within a single genome.

### Fine-tuning PlantCAD2 predict cross-species gene expression and protein abundance

To evaluate PlantCAD2’s ability to capture gene regulatory signals, we fine-tuned PlantCAD2 models using LoRA for two complementary tasks: gene expression and leaf translation (**Figure 6A, 6D**). Each task involved both classification (on/off status) and regression (absolute expression/translation level) objectives. For gene expression, we used promoter and terminator sequences (1024 bp each) as input; for translation, we used 500 bp upstream sequences. Following the same strategy as the accessible chromatin prediction task, we fine-tuned PlantCAD2 with LoRA and compared its performance with supervised PlantCAD2-S, CNN+LSTM, and AgroNT.

**Figure 6.**
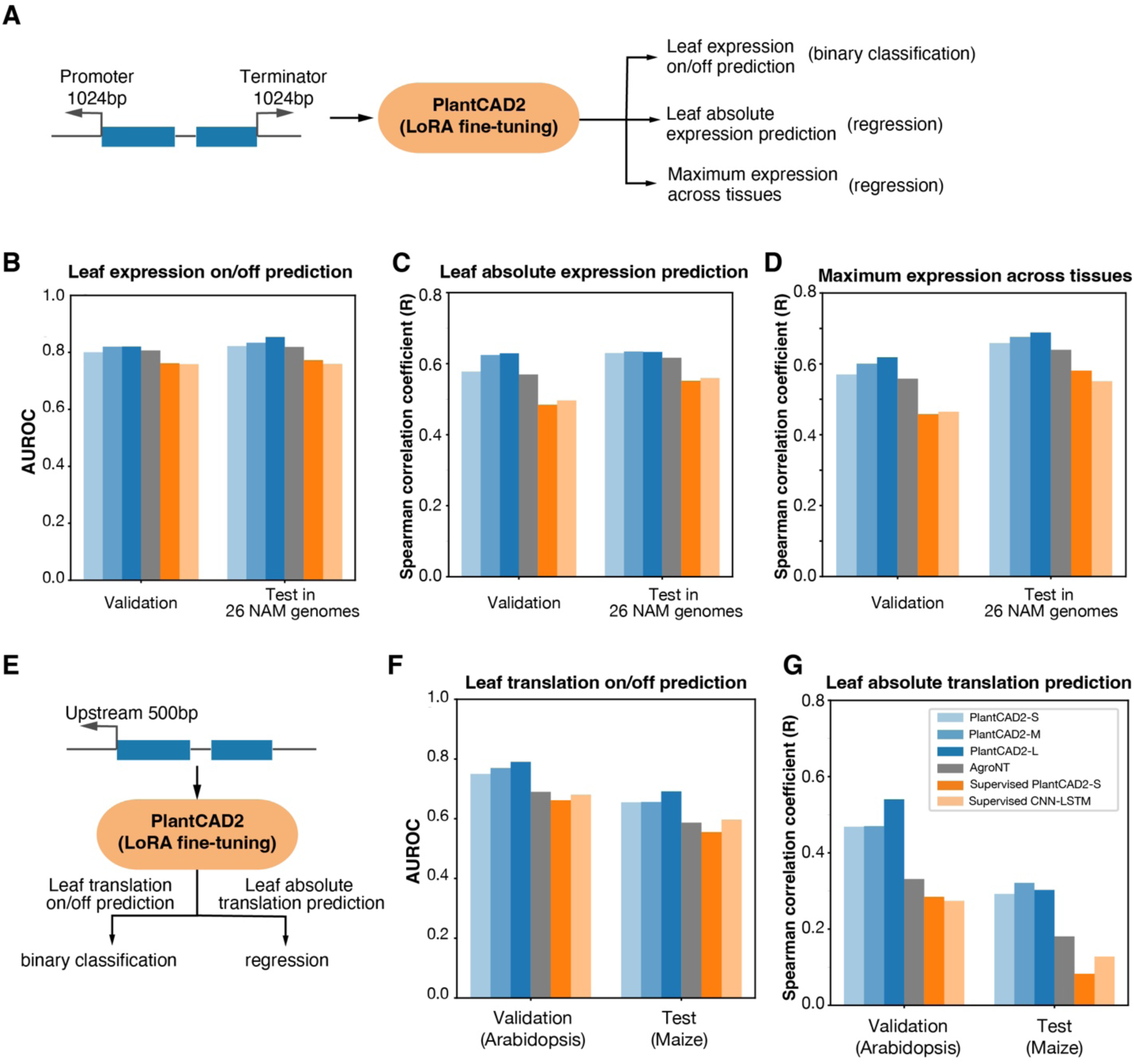
PlantCAD2 predicts gene expression and translation across species. **(A)** Gene expression prediction pipeline using promoter and terminator sequences (1024 bp each) for binary classification and regression tasks. **(B-D)** Cross-species gene expression performance on maize NAM population for binary on/off prediction (B), absolute expression levels (C) and maximum gene expression across tissues. **(D)** Translation prediction pipeline using 500 bp upstream sequences. **(E-G)** Translation prediction performance trained on Arabidopsis and tested cross-species on maize for binary on/off prediction (**F**) and absolute translation levels (**G**).

For cross-species gene expression modeling, we fine-tuned PlantCAD2 on a diverse panel of 15 plant species and evaluated predictions in 26 maize Nested Association Mapping (NAM) genomes ^71^ (Methods). We used three tasks to evaluate different models’ performance to predict gene expression: i) a binary classification task to predict if a gene is expressed or not expressed in leaf tissue; ii) a regression task to predict the absolute expression in leaf tissue; iii) a regression task to predict the maximum expression across tissues which represents a gene’s maximal transcriptional potential and avoids dilution from inactive tissues (**Figure 6A**). Across these tasks, PlantCAD2 consistently outperformed established baselines such as AgroNT and supervised CNN+LSTM. When testing in 26 NAM genomes, even the smallest PlantCAD2 (PlantCAD2-S, 88M parameters) outperformed the performance of the much larger AgroNT model (1B parameters) (**Figure 6B-6D**; **Supplemental Table 8**), demonstrating the efficiency of our foundation model framework. The largest model, PlantCAD2-L, achieved the best AUROC for binary leaf expression and the highest Spearman correlation for absolute expression prediction. To also rigorously assess predictive fit for regression tasks (ii and iii), we also reported Pearson correlation, R^2^, MAE (mean absolute error) and MSE (mean squared error) (**Supplemental Table 8**), observing consistent model performance rankings across these metrics. To further validate the model’s generalization capabilities beyond maize, we extended our evaluation to rice and tomato, where PlantCAD2 maintained the best cross-species performance (**Figure S11**).

Given that regulatory information also lies outside the proximal promoter ^38^, we evaluated the effect of varying input window sizes on gene expression prediction. Increasing the window from 1 kb to 4 kb both upstream of the transcription start site and downstream of the transcription stop site resulted in measurable improvements, raising the AUROC from 0.8221 to 0.8455 for binary leaf expression task and Spearman correlation from 0.6296 to 0.6455 for absolute expression prediction task on the NAM test set (**Figure S12**). To confirm that these improvements represent a true biological signal rather than stochastic variance, we performed repeated evaluations with different random seeds. The results exhibited high stability with minimal standard deviation (average SD = 0.0025 for binary leaf expression task and SD = 0.0032 for absolute expression prediction task; **Supplemental Table 9**; **Figure S12**), confirming that the performance gain from extended context—though numerically modest—is statistically significant and consistently reproducible. These improvements highlight the role of distal enhancers and long-range motifs in shaping expression. However, previous studies in both humans and plants have shown that current deep learning models lack the resolution to capture allele-specific effects ^71–73^. We therefore tested whether fine-tuning a foundation model could mitigate this limitation by evaluating per-orthogroup correlations within the NAM population, a comparison sensitive to allelic differences. Consistent with prior findings ^71–73^, only marginal improvements were observed for leaf absolute expression prediction, with the median Spearman correlation increasing from 0.112 (supervised CNN+LSTM) to 0.140 (PlantCAD2-S) (**Figure S13**). These small gains likely reflect both biological and data-related limitations of allele-specific modeling. Cis-regulatory elements such as transcription factor binding sites often exhibit high turnover rate ^74^, context-dependent, and effects driven by variants that are not explicitly represented in the training objective. In addition, robust allele-specific evaluation requires extensive data curation—such as accurate partitioning of cis and trans effects ^75,76^ and access to large-scale hybrid datasets with phased haplotypes or experimentally validated cis-regulatory variants. Such resources remain limited, and assembling them requires substantial experimental and computational curation, making it challenging for sequence-to-function models to learn and generalize allele-specific regulatory signatures. Together, these results suggest that achieving robust allele-specific expression prediction will require future models to incorporate explicit cis-variant modeling (e.g., VariantFormer ^77^) or fine-tuning on curated regulatory variant datasets ^75,78^.

Translation prediction was based on ribosome profiling (ribo-seq), a sequencing-based approach that estimates translation activity by mapping ribosome-protected mRNA fragments. Given high-quality ribo-seq datasets are scarce in plants, we restricted training to Arabidopsis and tested both within-species performance and cross-species transfer to maize. Interestingly, although supervised PlantCAD2-S (88M) is much larger than the CNN-LSTM (∼1.7M), the latter performed better, suggesting that large supervised models are prone to overfitting when trained on limited data (**Figure 6G**; **Supplemental Table 10**). By contrast, fine-tuned PlantCAD2 with parameter-efficient LoRA maintained robust performance without signs of overfitting. However, cross-species regression of absolute translation levels was less effective (**Figure 6G**), likely due to noise inherent in current ribo-seq–based quantitative estimates. We anticipate that as more high-quality cross-species ribo-seq datasets become available, fine-tuning will demonstrate significantly improved robustness for quantitative prediction. Despite this limitation, we were encouraged by the strong transfer observed for binary classification of leaf translation. We next tested whether gene expression could similarly be transferred from Arabidopsis to maize. Using a separate Arabidopsis gene expression dataset ^79^ with 1,024 bp upstream and downstream sequences, we found that direct transfer performed poorly (AUROC: 0.786 in Arabidopsis vs. 0.631 in maize) compared to protein abundance prediction (0.790 in Arabidopsis vs. 0.692 in maize) (**Figure S14**). This contrast may highlight a fundamental difference between regulatory layers: translational control appears to be more evolutionarily conserved than transcriptional regulation. Consistent with evidence that protein abundance is under stronger selective constraint than transcript levels ^63^, these results explain why translation prediction transfers effectively across species, whereas accurate gene expression prediction requires phylogenetically diverse training data.

## Discussion

In this work, we present PlantCAD2, an extended-context window DNA language model that substantially advances the sequence-to-function modeling in plant genomics. Building on the foundation laid by PlantCAD, PlantCAD2 features a model architecture that is three times larger, a 16-fold longer context window (8,192bp vs. 512bp), and a pre-training dataset that is evenly distributed across angiosperm orders to better capture phylogenetic diversity. Through comprehensive zero-shot and fine-tuned evaluations, we demonstrate that PlantCAD2 not only exhibits strong cross-species generalization, but also achieves superior performance across a wide range of sequence-to-function tasks, including evolutionary conservation prediction, functional important junction sites prediction involved in both transcription and translation, variant (including indels) effect estimation, cis-regulatory activity, gene expression, and protein translation.

A key design consideration for PlantCAD2 is the careful curation of 65 angiosperm species to maximize phylogenetic diversity while minimizing data bias. Rather than using all publicly available plant genomes, we selected a single representative genome per genus. This balanced strategy avoids two major issues. First, pre-training on all available genomes would dramatically increase computational cost. As shown by scaling laws in large language models, increasing dataset size alone while keeping model size fixed leads to diminishing returns unless model capacity and training compute are expanded proportionally ^80^. In practice, simply adding more genomes would not necessarily improve performance unless the model were enlarged substantially, which would reduce its efficiency and complicate deployment. Second, public genome resources are heavily skewed toward a few lineages such as grasses and Brassicaceae^81^. Such skewed distributions have been widely documented across NLP ^82,83^ and protein language models ^84^. These studies consistently show that large models tend to learn disproportionately from overrepresented groups in the training data, resulting in biased representations and reduced generalization to underrepresented categories. This highlights the importance of deliberate data curation to ensure balanced phylogenetic coverage and to prevent the model from overfitting to a few dominant plant lineages.

Furthermore, we emphasize that phylogenetic diversity is often more important than the absolute number of species used for pre-training ^35^. This has been observed in models such as NTv2 ^25^, which achieve strong performance with a few hundred phylogenetically diverse species, outperforming models trained on much larger but with 1,000 human genomes. This distinction is particularly critical when comparing PlantCAD2 to models trained on broader evolutionary scales. While generalist models such as Evo2 and NTv3 ^85^ incorporate genomes spanning the entire tree of life, recent results suggest that this extreme phylogenetic breadth may come at the cost of lineage-specific representation quality. Notably, NTv3 achieves strong performance across diverse tasks when fine-tuned with supervised data, yet its zero-shot performance on all 12 plant-specific benchmarks is substantially lower than PlantCAD2 (**Figure S16**). This pattern parallels the Evo2 results reported above and highlights an important distinction: fine-tuning can compensate for weaker lineage-specific representations through task-specific parameter updates, but zero-shot performance — which directly reflects what a model has internalized during pre-training — provides a more direct measure of representation quality for a given clade. By strictly focusing on angiosperms—which diverged around 160 million years ago and share a coherent regulatory grammar—PlantCAD2 captures generalizable constraints specific to flowering plants without the "noise" of unrelated genomic architectures (e.g., prokaryotic or mammalian regulatory logic). This targeted approach allows PlantCAD2 to achieve superior performance on plant-specific tasks while maintaining computational efficiency.

The choice of an 8,192 bp context window reflects a deliberate balance between biological resolution and computational feasibility. Although distal regulatory interactions in plants can extend over tens of kilobases ^38^, our survey of the 65 training genomes shows that the median gene length is 2,422 bp (**Figure S15**), therefore, an 8 kb window is sufficient to encompass most gene bodies together with their core regulatory regions. Regions far beyond this range are frequently dominated by transposable elements rather than functional cis-regulatory modules ^86^, reducing their contribution to prediction accuracy. From a technical perspective, even with efficient state-space architectures such as Mamba, both computation time and GPU memory increase approximately linearly with context length ^37,42^, making substantially longer windows (e.g., 16–32 kb) impractical for many users and typical hardware settings. Importantly, PlantCAD2 enables flexible chunking strategies that allow the model to integrate distal information across adjacent windows when needed. Together, these considerations make 8 kb a pragmatic choice that maximizes biological utility while maintaining accessibility and efficiency.

PlantCAD2 represents a significant step toward a foundational model for plant genomics. Rather than building task-specific models for each application, it enables unified modeling of sequence-to-function relationships that can be efficiently adapted across cell types, tissues, and species. Fine-tuning PlantCAD2 enables accurate de novo genome annotation ^87^, further expanding its utility for genome interpretation. This paradigm shift opens new opportunities to integrate deep learning into practical breeding applications. For example, PlantCAD2 could assist in prioritizing causal variants in GWAS studies, interpreting SVs in noncoding regions, or guiding sequence design for synthetic promoters with desired expression patterns. Its ability to transfer knowledge across evolutionarily distant species further enhances its utility for crop improvement, particularly in non-model organisms where high-quality training data are limited but genomic sequences are available.

Despite its advances, PlantCAD2 also presents new challenges. First, its large model size may limit deployment in GPU-constrained environments. To directly assess the lower bound of model size, we additionally trained and evaluated a substantially smaller PlantCAD2 model (∼20M parameters) using the same pre-training data and zero-shot evaluation protocols. In contrast to the PlantCAD2-S, this lightweight model showed a pronounced drop in performance across most tasks (**Supplementary Table 11**). For example, in predicting evolutionary conservation within Andropogoneae, the AUROC decreased from 0.725 (PlantCAD2-L) to 0.593 (20M model), indicating a clear capacity threshold below which biologically meaningful sequence representations cannot be reliably learned. These results suggest that the smallest released model (88M parameters) already represents a near-minimal configuration that balances predictive accuracy and computational efficiency, and that further reductions in parameter count will likely require model compression or distillation strategies rather than naive scaling. Second, there are still technical barriers that may limit accessibility to wet-lab biologists or breeders. Building intuitive interfaces, pretrained APIs, and end-to-end pipelines will be crucial to broaden the use of PlantCAD2 in broader plant science communities, and these are obvious areas for future development. Third, while the 8,192-bp context window allows PlantCAD2 to model distal regulatory elements, further extending this capability would be valuable for capturing long-range interactions such as enhancer–promoter loops. For example, in maize, the teosinte branched 1 (*tb1*) enhancer ^88,89^ and Vegetative to generative transition 1 (*Vgt1*) ^90^ are located approximately 70kb and 60kb upstream of their target genes, respectively. However, capturing such interactions will likely require novel tokenization or compression strategies that can represent long, repetitive sequences without sacrificing resolution.

Looking forward, future directions include combining PlantCAD2 with multi-modal data such as DNA methylation and chromatin states will provide trans-factors to the genome. In addition, diffusion-based sequence generation models could also be promising coupled with synthetic biology. Ultimately, we envision PlantCAD2 and its successors as key building blocks for a sequence-to-function foundation model capable of enabling predictive genomics and rational genome design in diverse plant species.

## Methods

### Preparing pre-training genomes

A total of 65 genomes were selected for pre-training from the Phytozome database. To ensure taxonomic relevance and minimize redundancy, we applied a series of manual filtering steps. First, non-angiosperm species were excluded. For each remaining species, we retained only the most recent genome assembly version. In cases where two haplotypes were available for a species, we selected the haplotype with the higher N50 value; if N50 values were comparable, we retained the assembly with fewer scaffolds to prioritize less fragmented genomes. Taxonomic information, including order, family, and genus, was appended to each genome to facilitate downstream analyses, and their relationships were visualized using a published time-callibrated phylogeny ^91^. For each selected genome, we extracted genomic sequences centered on each annotated gene, extending 5 kilobases (kb) upstream and 5 kb downstream from the gene center. These ±5 kb gene-centered regions were then segmented into overlapping windows of 8,192 bp with a step size of 4,096 bp, ensuring comprehensive coverage of regulatory and genic features while maintaining continuity across sequence boundaries. These windows served as input sequences for model pre-training.

### PlantCAD2 model architecture and pre-training

PlantCAD2 builds upon the Caduceus architecture ^26^ used in PlantCAD ^27^, retaining its key design principles while incorporating architectural improvements. Like PlantCAD1, PlantCAD2 maintains three core features: (1) bidirectional sequence processing, where sequences are processed both forward and reverse with outputs summed together; (2) reverse-complement (RC) equivariance, ensuring the model commutes with RC operations; and (3) parameter-efficient bidirectional implementation through shared linear projections between forward and reverse passes.

The primary architectural improvement in PlantCAD2 is the replacement of Mamba1 blocks ^37^ with Mamba2 blocks ^42^. Mamba2 introduces a structured state space duality that recasts the selective state space computation into an equivalent convolutional form using structured matrices, improving parallelism and hardware efficiency. This dual representation enables significantly faster training (up to 2–4× in some scenarios) while retaining the input-dependent selection mechanism that allows the model to dynamically modulate state updates based on sequence content. These advances allow PlantCAD2 to efficiently handle 8,192 bp sequences with linear computational complexity.

For the pre-training of PlantCAD2, each model was trained for 240,000 steps using a Decoupled AdamW optimizer ^92^ with the global batch size of 2,048. The learning rate is 2E-4 with a cosine decay scheduler, and 6% of the training duration was dedicated to warm up. The learning rate decayed to 4E-6 by the end of training. The default BERT ^34^ masking recipe was used with a masking probability of 15%. For each masked token: i) there is an 80% probability it will be replaced by a special token ([MASK]), ii) a 10% probability it will be replaced by a random token, and iii) a 10% probability it will remain unchanged. Unless otherwise specified, all models were trained using a sequence length of 8192 base pairs. A weight decay of 1E-5 was applied throughout the training process. Pre-training of the 676M-parameter PlantCAD2-L model required approximately 28 days on 64 NVIDIA H100 GPUs, while the 88M-parameter model completed training in approximately 3 days under the same hardware configuration.

### Repeat annotation and loss re-weighting

To mitigate the influence of highly repetitive sequences during pre-training, we explicitly down-weighted the contribution of repetitive regions to the masked language modeling loss. This loss has been found in other DNA models ^22,27^ to improve performance on downstream tasks and better calibrate likelihoods between repetitive and nonrepetitive DNA. We employed a de-novo pipeline to annotate highly repetitive sequences ^93^ (https://github.com/baoxingsong/dCNS), then repetitive sequences are down-weighted during pre-training as demonstrated important in previous studies ^22^. Then the loss is calculated as:

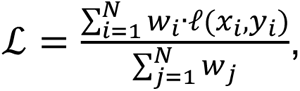

Where *w_i_* represents the weight of i-th nucleotide,

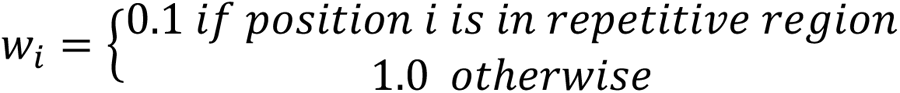

### Evolutionary constraint prediction using the zero-shot strategy

To evaluate the extent to which PlantCAD2 captures evolutionary conservation signals, a zero-shot strategy was applied to predict constrained genomic regions. Two independent tasks were used. The first task focused on *Sorghum bicolor*, using conservation estimates from the Andropogoneae tribe, a large clade of approximately 1,200 grass species that descended from a common ancestor approximately 18 million years ago ^94^. To generate conservation labels, 34 high-quality genomes ^95^ were aligned to the Sorghum bicolor reference genome using AnchorWave ^96^. Per-base conservation was quantified using alignment identity scores across all species. Sites with high-quality coverage (i.e., aligned in at least 34 out of 35 species) were retained for analysis. Among these, positions with an identity score ≥34 were labeled as conserved, while those with identity scores <15 were labeled as neutral. Sites with intermediate identity scores or insufficient coverage were excluded from evaluation to ensure high-confidence labels ^27^.

The second task utilized conservation scores derived from multiple sequence alignments (MSAs) of orthologous coding sequences from 325 Poaceae genomes, a high-quality subset of the recently published set of 727 genomes. Using *Pharus latifolius* as an outgroup species, gap columns were removed prior to conservation estimates. Using PHAST ^97^, PhyloP scores were calculated per site based on a neutral model derived from fourfold degenerate sites and “LRT” methods with the mode ”CONACC”. Sites with phyloP scores above 5 were classified as conserved, while those below 1.5 were considered neutral. Sites with intermediate scores were excluded to maintain label clarity. For TIS sites, we retained all 36,668 (26,653 conserved vs 10,015 less conserved) sites given their biological importance. For non-TIS sites, we downsampled to 183,687 sites (103,369 conserved versus 80,318 neutral) for computational efficiency while maintaining the conserved-to-neutral ratio.

For both tasks, the evaluated site was centered (i.e., 4096^th^ of 8,192bp) within an input sequence, and the reference base at that position was masked. The model’s predicted likelihood of the reference allele was extracted and used as the zero-shot conservation score. Higher likelihoods were hypothesized to reflect stronger conservation. Model performance was assessed using AUROC, comparing scores between conserved and neutral sites.

### Core and non-core gene classification using the zero-shot strategy

To assess the ability of PlantCAD2 to distinguish between core and non-core genes in a population, a zero-shot strategy was applied to classify within species conservation of genes in maize and tomato. For maize, the pangene table derived from 26 Nested Association Mapping genomes ^98^ was used. Core genes were defined as those present in all 26 NAM genomes, whereas non-core genes included both dispensable genes (present in 2-23 genomes) and private genes (present in only one genome). For genes with multiple transcripts, the canonical transcript specified in the annotation was used. For tomato, the pan-genome dataset assembled from 586 high-quality genomes ^99^ was used. Genes present in all 586 accessions were defined as core genes, and non-core genes consist of dispensable (present in 6-580 accessions) and private (present in less than 5 accessions). The longest transcript was selected to represent each gene across all analyses.

To quantify the model’s prediction at each functional junction, a masked motif accuracy score was defined. For example, to evaluate translation initiation sites, the canonical ATG start codon was masked, and the model’s predicted likelihoods for the three masked nucleotides were extracted and averaged. A similar approach was applied to other junction types, including translation termination sites (TAA, TAG, TGA), splice donor sites (GT), and splice acceptor sites (AG), by masking the corresponding motifs and calculating average token likelihoods.

To evaluate performance, core genes were treated as positive and non-core genes as negative, and AUROC was calculated based on the masked motif accuracy scores for each gene.

### Accessible chromatin region prediction

To evaluate the capability of PlantCAD2 to capture regulatory sequence features, fine-tuning experiments were performed using Low-Rank Adaptation (LoRA) ^68^ on two accessible chromatin prediction tasks: (1) cross-species accessible regions (ACRs) prediction, and (2) cell-type-specific ACR prediction.

For the cross-species task, ATAC-seq peak regions from 12 plant species ^67^ were downloaded from NCBI (https://www.ncbi.nlm.nih.gov/geo/query/acc.cgi?acc=GSE128434). We followed the data processing pipeline as described by Wrightsman et al ^100^. For each species, peak regions were processed by extracting the midpoint of each peak and symmetrically extending it by half the target input window size (300bp, 600bp and 1,000bp) in both directions to generate positive observations. To reflect the real-world scenario in which most of the genome is inaccessible, the rest of the genome was used as negative examples, ensuring no overlap with known peaks.

For the cell-type-specific task, we used the single-cell ATAC-Seq ^70^ and used a similar preprocessing pipeline, but each genomic region could be associated with accessibility across 92 cell types. As such, the task was framed as a multi-label classification problem, where each region was assigned a binary accessibility label for each of the 92 cell types based on its overlap with experimentally identified peaks.

### Gene expression prediction in leaf

To evaluate the models’ ability to predict gene expression, we designed two tasks: (1) leaf absolute expression and (2) leaf on/off expression classification. The training dataset was derived from 15 Andropogoneae species ^71^. For validation, we held out two species closest to *Zea mays*—*Tripsacum zopilotense* and *Zea diploperennis*—both members of the Tripsacinae subtribe, which diverged from maize approximately 0.6 to 4 million years ago. This setup enabled evaluation of the models’ cross-species generalization to closely related taxa. For the leaf absolute expression task, the log_10_TPM values were used as regression targets during fine-tuning. For the on/off expression task, genes with TPM > 1 were labeled as expressed (positive), and those with TPM ≤ 1 were considered non-expressed (negative). This setup enabled evaluation of the models’ ability to generalize expression predictions across closely related species within the clade.

### Leaf protein abundance prediction task

To evaluate the models’ ability to predict protein abundance, we designed two tasks analogous to the gene expression analysis: absolute abundance and on/off classification. Ribo-Seq data were obtained from Arabidopsis ^101^ and *Z. mays* ^102^. Raw reads were downloaded from NCBI, and Trimmomatic ^103^ was used to trim adapters and filter low-quality reads. Cleaned reads were first aligned to rRNA reference sequences using Bowtie ^104^ to remove contaminating rRNA. The remaining reads were then mapped to the reference genomes of Arabidopsis and maize using STAR ^105^. Gene-level translation abundance was quantified using StringTie ^106^ based on uniquely mapped reads. We then designed two tasks analogous to the expression prediction setup: (1) absolute protein abundance, where the log10-transformed Ribo-Seq expression values were used as regression targets, and (2) on/off classification, where genes with TPM > 1 were considered expressed (positive) and those with TPM ≤ 1 were labeled as non-expressed (negative).

### Fine-tuning PlantCAD2

To adapt the pre-trained PlantCAD2 model to downstream tasks, we employed Low-Rank Adaptation (LoRA) ^68^, a parameter-efficient fine-tuning strategy that inserts trainable low-rank matrices into the attention layers of the transformer. This approach enables effective adaptation while keeping the vast majority of model parameters frozen. Fine-tuning was performed using the PEFT library ^107^ with LoRA rank = 8, α = 32, and dropout = 0.1, targeting the "x_proj", "in_proj", and "out_proj" modules. Models were trained using the Hugging Face Trainer with a learning rate of 1e−4, a global batch size of 128, and one training epoch. BF16 precision and linear learning rate scheduling with 50 warm-up steps were used. Over 98% of the model parameters remained frozen, enabling efficient and scalable fine-tuning across tasks. All tasks were fine-tuned for a single epoch without hyperparameter tuning to ensure stability and consistency across experiments. Fine-tuning objectives for all models were a binary cross entropy loss for classification tasks and a mean squared error loss for regression tasks.

### Fine-tuning AgroNT

To directly compare the performance of fine-tuned PlantCAD2 with AgroNT ^24^, we applied the same parameter-efficient fine-tuning strategy using LoRA. All LoRA hyperparameters were kept consistent with those used for PlantCAD2, including rank = 8, α = 32, and dropout = 0.1. For AgroNT, LoRA adapters were inserted into the "query" and "key" projection layers of the transformer ^43^, reflecting its architecture. This setup ensured a fair comparison between models under matched fine-tuning conditions.

### Supervised CNN + LSTM baseline

To benchmark against traditional supervised models, we implemented a CNN+LSTM architecture based on DanQ ^108^, a widely used hybrid model for DNA sequence classification. For each task, the model was trained from scratch using one-hot encoded sequences. We used the Adam optimizer with a learning rate of 0.01, a batch size of 2,048, and trained for up to 200 epochs, with early stopping after 20 epochs without validation improvement.

### Supervised PlantCAD2 baseline

To assess the contribution of pretraining, we trained a small PlantCAD2 model from scratch for each downstream task. This supervised baseline used the same architecture and hyperparameters as the fine-tuned version but was initialized without pretrained weights—by loading only the Hugging Face model configuration.

### Zero-shot evaluation of PlantCAD2, PlantCAD, GPN and NTv3 models

All three models were pre-trained with masked language modeling. For PlantCAD, we used the largest available model "kuleshov-group/PlantCaduceus_l32" for evaluation. For GPN, we used "songlab/gpn-brassicales". Due to the 512 bp context window limitation of both PlantCAD and GPN, we cropped input sequences to 512 bp centered on the target position. For NTv3, we used the largest pre-trained version (InstaDeepAI/NTv3_650M_pre) for evaluation. All other evaluation configurations remained identical to those used for PlantCAD2.

### Zero-shot evaluation of Evo2 model

All zero-shot tasks were also benchmarked using the Evo2 ^19^ model (“evo2_7b”) for comparison. Since Evo2 is autoregressive (predicting the next token rather than masked tokens), masked token accuracy could not be directly computed. Therefore, for the evolutionary constraint task, we fed the full input sequence into the model and extracted the likelihood of the reference allele at the target site as the conservation score. To ensure a fair comparison, we used an 8,192 bp context window for Evo2, matching the input length used for PlantCAD2 evaluations. The same approach was applied for benchmarking structural variants.

For the masked motif accuracy task, we evaluated Evo2 using two configurations to compensate for its unidirectional architecture: (1) forward sequences (Evo2-fwd), where the model uses upstream context to predict the junction—for example, for TIS prediction, we used the 4,094 bp upstream of the TIS as a prompt for Evo2 to generate the next three tokens; and (2) reverse-complement sequences (Evo2-rc), where the model uses downstream context (reverse complemented) to predict the junction in the opposite direction.

## Code availability

All pre-trained models, datasets, and benchmark tasks are available at https://huggingface.co/collections/kuleshov-group/plantcad2-67e437e241a382671371a572.

Fine-tuning pipelines and code are available at https://github.com/plantcad/plantcad.

## Acknowledgements

This work is funded by a cooperative agreement from US Department of Agriculture Agricultural Research Service to Cornell University (MCR), NSF grant (#2240888), NSF CAREER grant (#2145577), NIH Maximizing Investigators’ Research Awards (#1R35GM151243-01 and 5R35GM151348). SWe thank Travis Wrightsman (Cornell University) for sharing gene expression tasks, Michelle C. Stitzer (Cornell University) for plotting the tree, Hai Wang (China Agricultural University) for sharing max expression data of 17 species, Yaoyao Wu (Nanjing Agricultural University) for providing conserved and neutral SNP datasets, Sara Miller (Cornell University) for helpful comments, and all members of Cornell’s Institute of Genomic Diversity (MCR & ESB) for helpful discussions. We would also like to thank the SCINet project, Texas Advanced Computing Center at The University of Texas at Austin, and MosaicML for providing compute resources for pretraining and fine-tuning experiments.

## Author contributions

J.Z., A.G., V.K., and E.S.B. designed research; J.Z., A.G., S.-K.H., Z.-Y.L., S.-P.C., E.M., E.C., B.C., A.B., M.C.R., M.P., V.K., and E.S.B. performed research and analyses; J.Z., E.M., S.-K.H., M.P., and E.S.B. wrote the manuscript with all other authors’ suggestions and comments.

## Supplemental Information

**Supplemental Table 1.** Pretraining species and masked language modeling performance across 65 angiosperm genomes

**Supplemental Table 2.** Cross-species evolutionary conservation prediction performance

**Supplemental Table 3.** Masked motif prediction accuracy for transcriptional and translational junction sites

**Supplemental Table 4.** Zero-shot structural variant impact prediction performance

**Supplemental Table 5.** Cross-species chromatin accessibility prediction trained on Arabidopsis

**Supplemental Table 6.** Multi-species chromatin accessibility prediction performance

**Supplemental Table 7.** Cell-type-specific chromatin accessibility prediction in maize

**Supplemental Table 8.** Gene expression prediction performance across specie

**Supplemental Table 9.** Gene expression prediction performance with five random seeds

**Supplemental Table 10.** Translation prediction performance across species

**Supplemental Table 11.** The zero-shot performance of the tiny PlantCAD2 model with 20M parameters

