## Supplemental Figures for "PlantCAD2: a DNA foundation model for interpreting genomes across flowering plants"

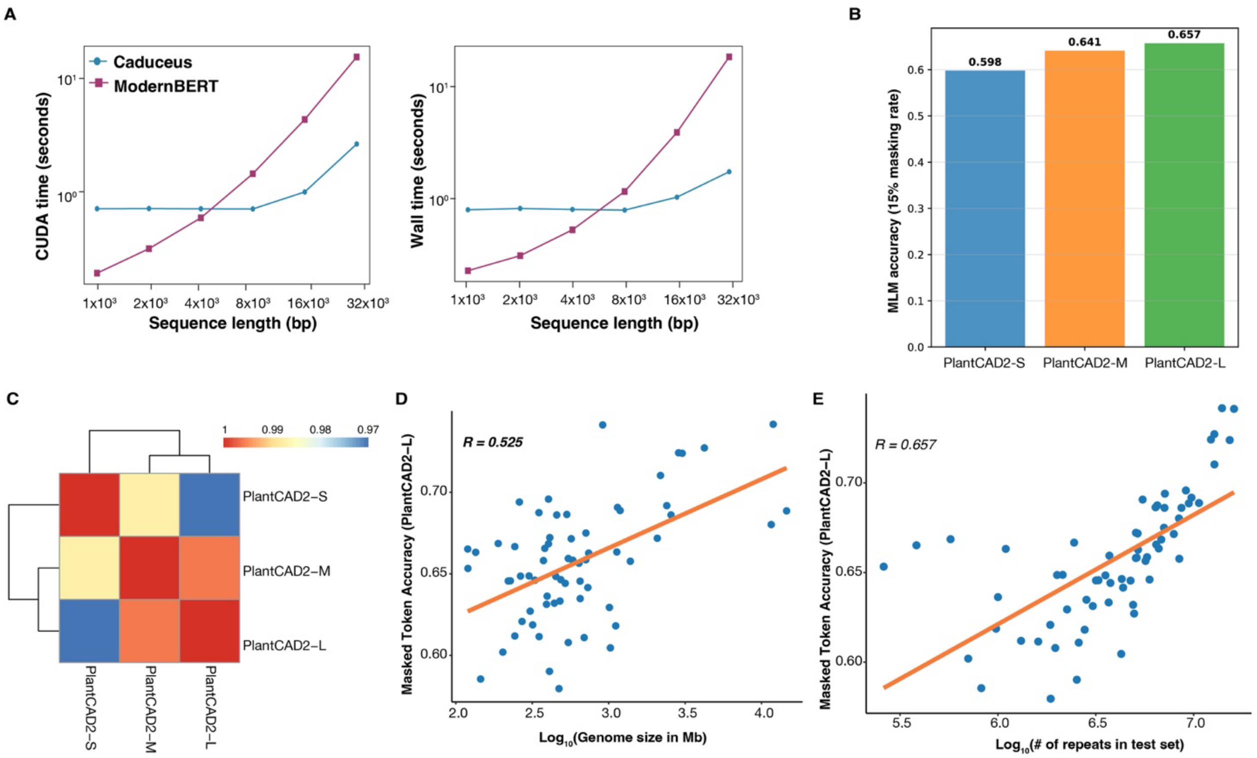


**Figure S1. PlantCAD2 model performance across different scales and species. (A)** Inference speed benchmarking shows that the Caduceus architecture exhibits significantly faster CUDA and wall-clock runtime than ModernBERT across increasing sequence lengths. **(B)** Masked language modeling accuracy across the three models, evaluated by randomly masking 15% of nucleotides per sequence, showing improved performance with increasing model size. **(C)** Correlation matrix of per-species masked language modeling accuracy of three PlantCAD2 models. **(D)** Relationship between genome size and masked language modeling accuracy for PlantCAD2-L. **(E)** Relationship between the number of repetitive sequences in the test set and masked language modeling accuracy.

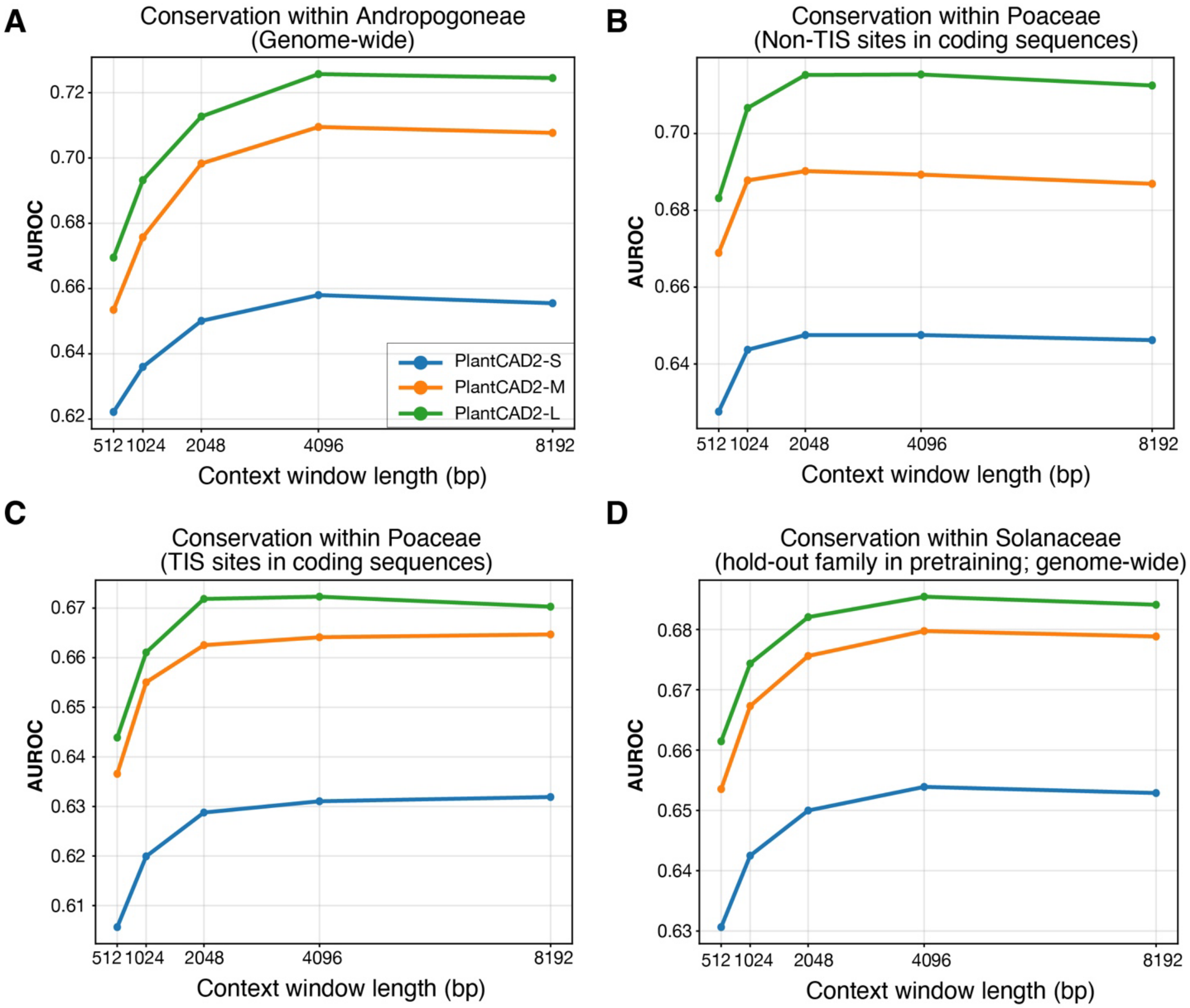


**Figure S2. Effect of context window length on AUROC performance.** Evaluation across three PlantCAD2 models for conservation within Andropogoneae **(A)**, Poaceae non-TIS **(B)**, Poaceae TIS **(C)**, and Solanaceae **(D)**.

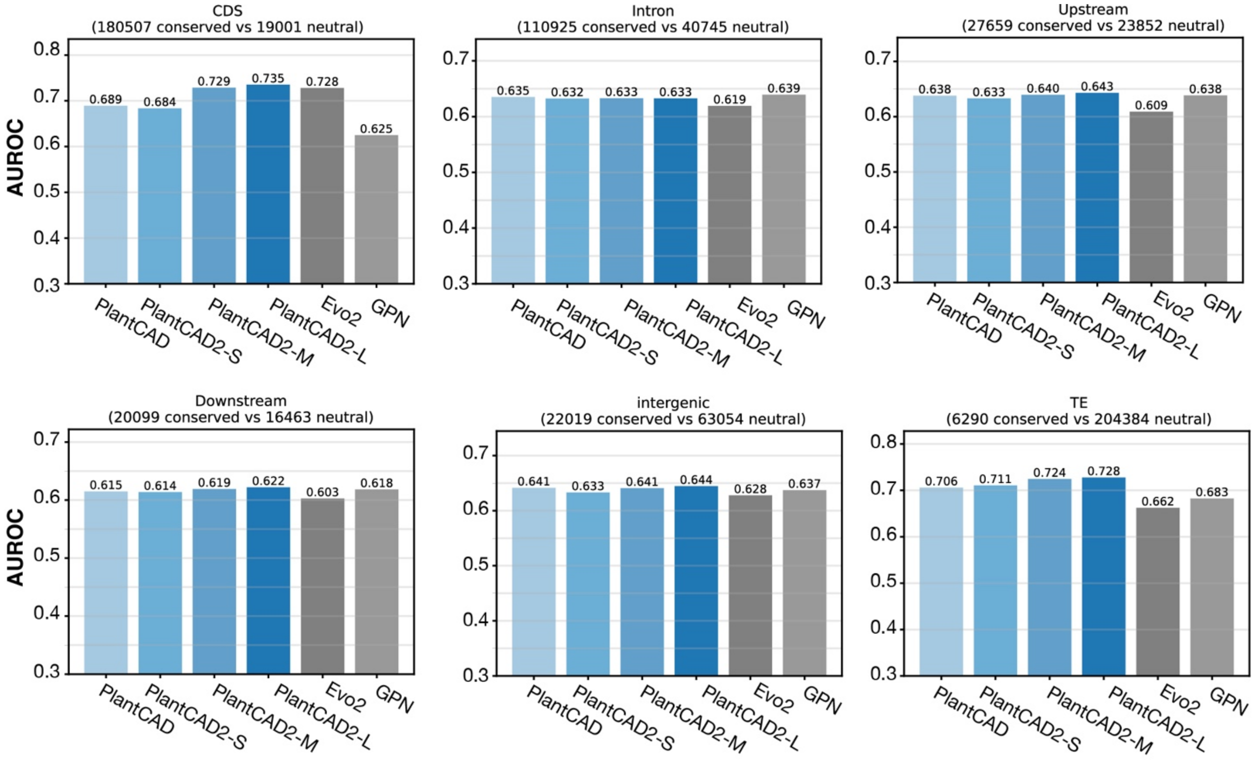


**Figure S3. AUROC of different genomic contexts for conservation prediction in Solanaceae.**

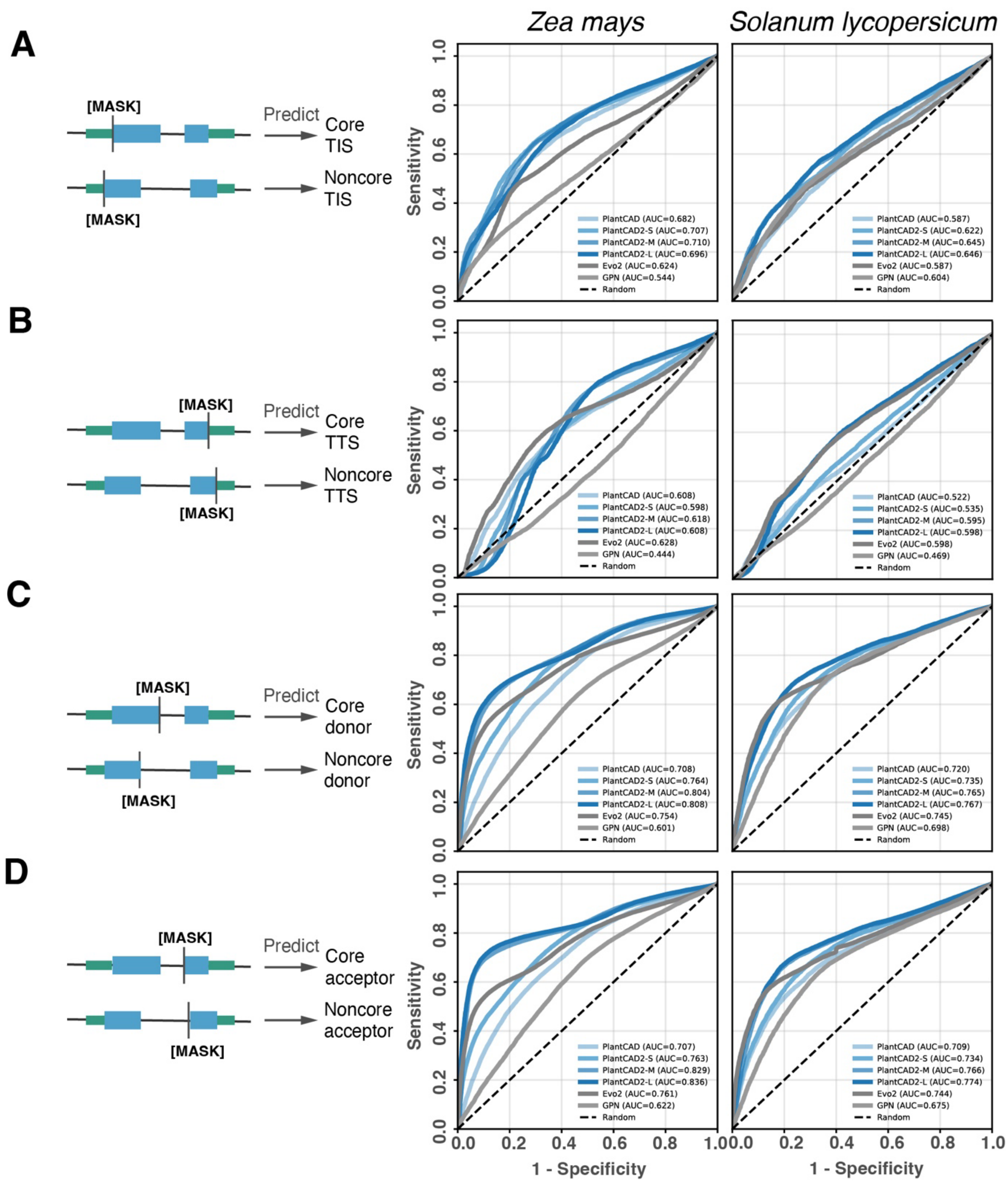


**Figure S4. PlantCAD2 distinguishes core from non-core junctions with zero-shot strategy.** ROC curves showing binary classification performance for distinguishing evolutionarily constrained core junctions from rapidly evolving non-core junctions using model log-likelihood scores at junction sites. Left panels show maize (included in pre-training), right panels show tomato (excluded from training). **(A)** Translation initiation sites. **(B)** Translation termination sites. **(C)** Splice donor sites. **(D)** Splice acceptor sites.

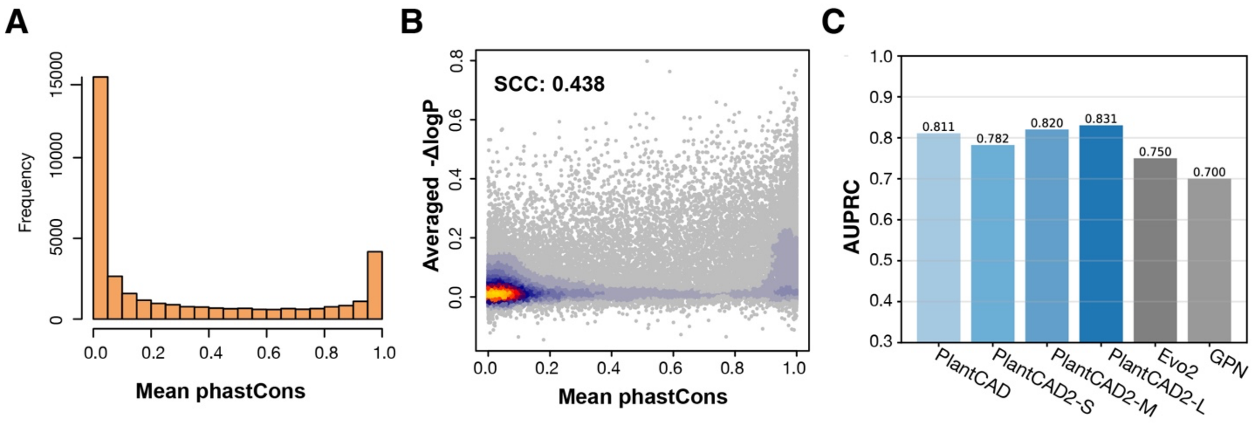


**Figure S5. PlantCAD2 predicts functional impact of structural variants using zero-shot strategy.** (A) Distribution of phastCons. (B) Scatter plot showing the positive correlation between PlantCAD2's -ΔlogP scores and phyloP-based conservation scores. (C) AUROC performance distinguishes highly conserved from less conserved deletions.

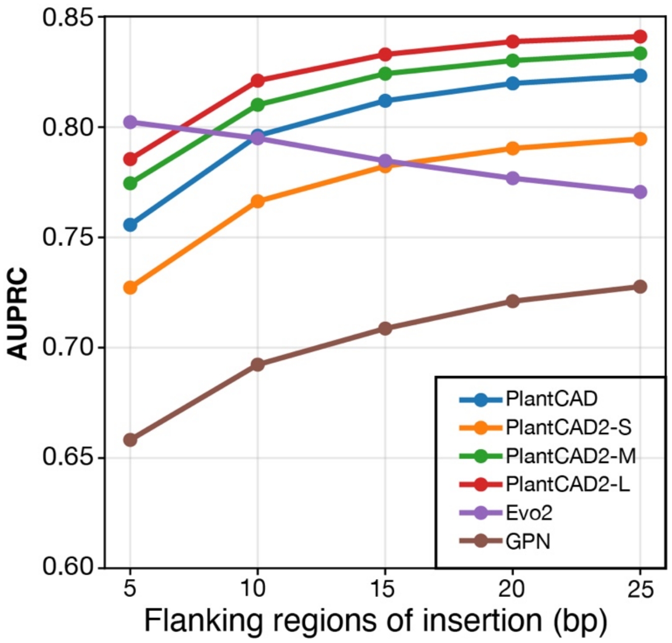


**Figure S6. Effect of flanking sequence length on deletion classification performance.** AUROC performance for distinguishing conserved from less conserved deletions by flanking sequence length.

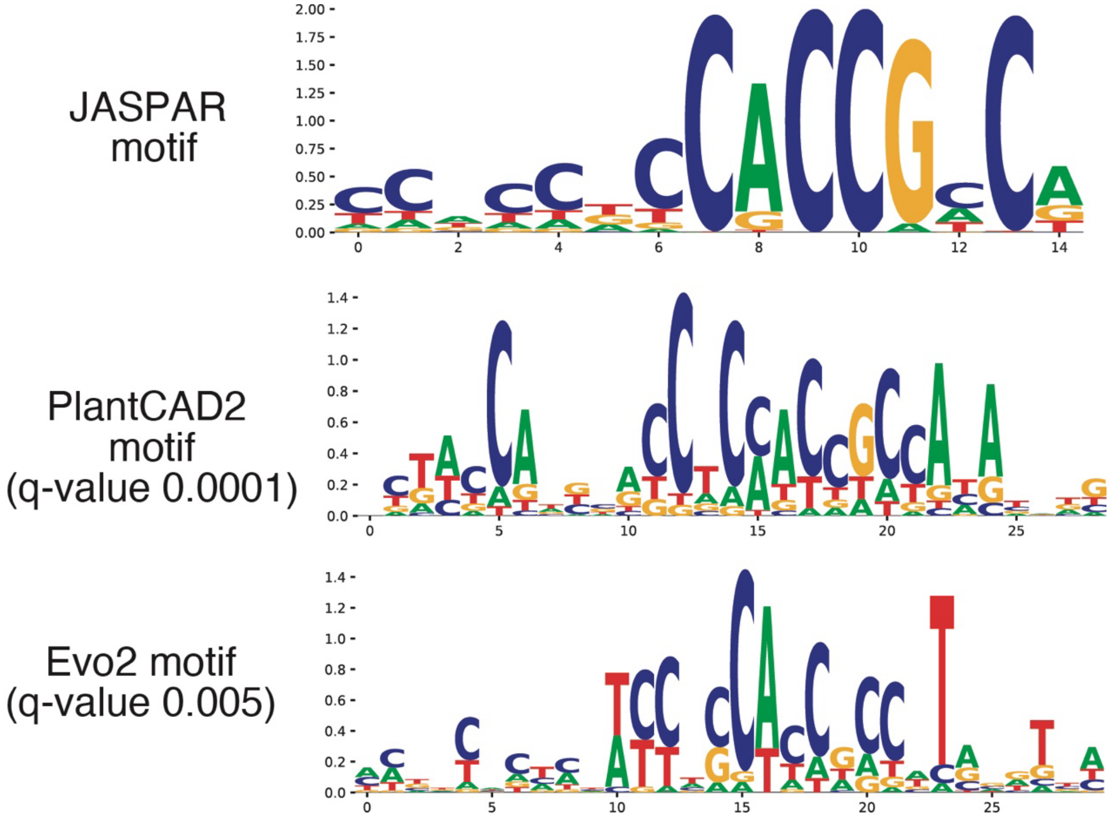


**Figure S7. Motifs generated by TF-MoDISco from PlantCAD2 and Evo2 high-confidence predictions for DREB2 (AP2/ERF family).**

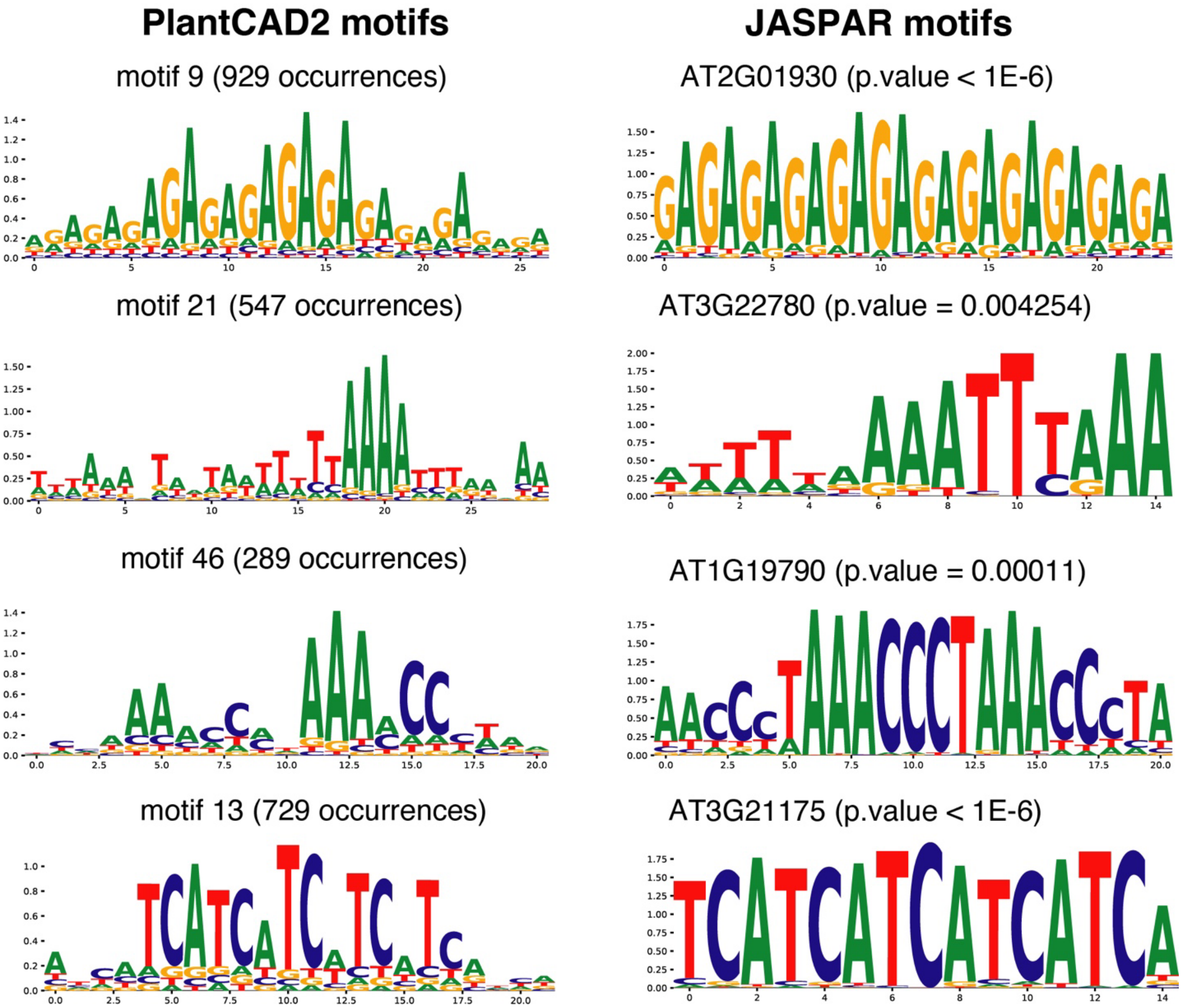


**Figure S8. Example motifs generated by TF-MoDISco from PlantCAD2 high-confidence predictions in Arabidopsis promoter regions.**

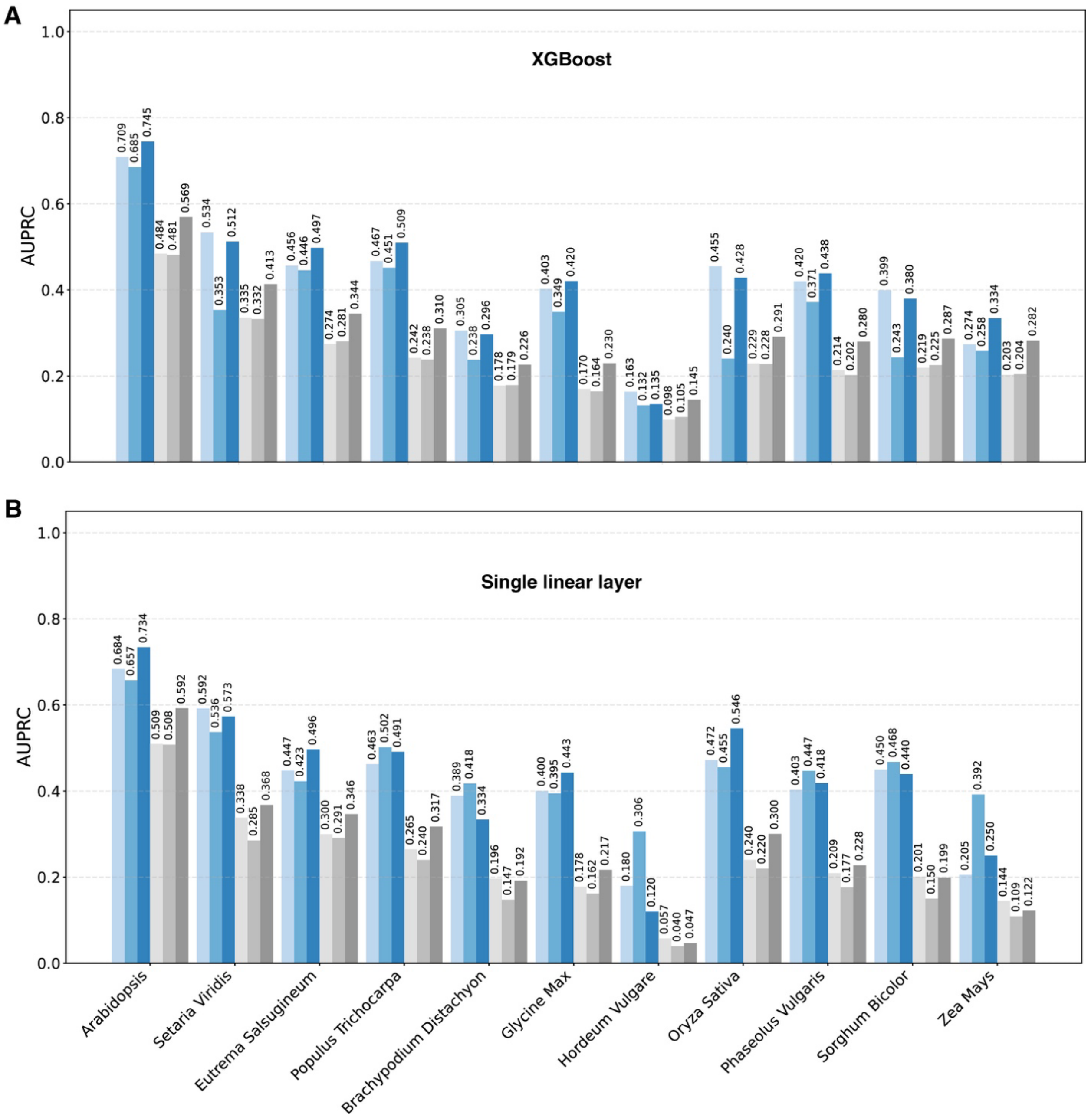


**Figure S9. Accessible region prediction task comparison with Evo2 using frozen embeddings. (A)** XGBoost classifier**, (B)** neural network with single linear layer.

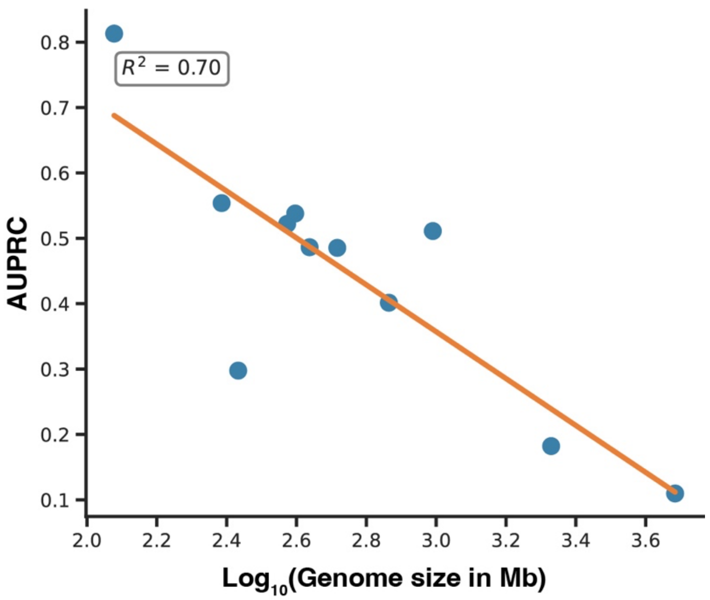


**Figure S10.** Relationship between genome size and accessibility prediction performance.

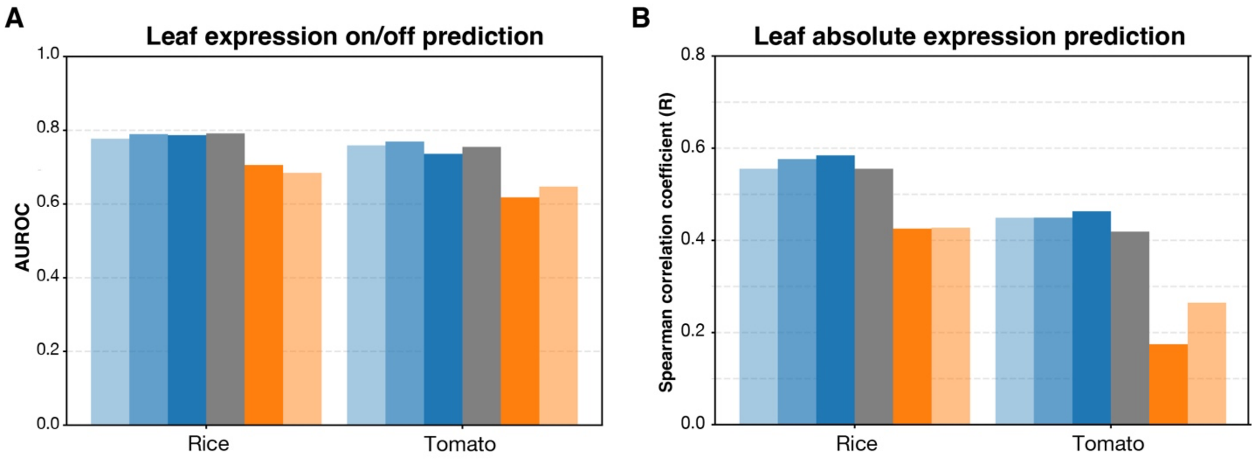


**Figure S11. Gene expression prediction performance in rice and tomato. (A)** Leaf expression on/off prediction**, (B)** leaf absolute expression prediction.

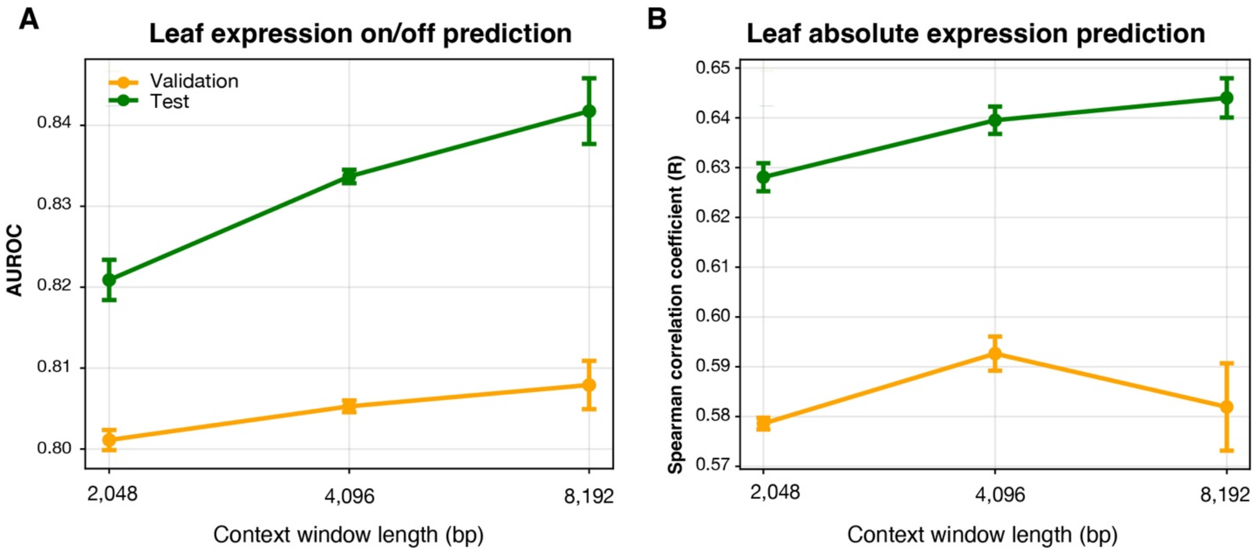


**Figure S12. Effect of context window length on gene expression prediction performance.** Performance improvement with increasing context window size for binary leaf expression classification **(A)** and absolute expression regression **(B)** on the maize NAM dataset. Error bars represent five replicates with different random seeds.

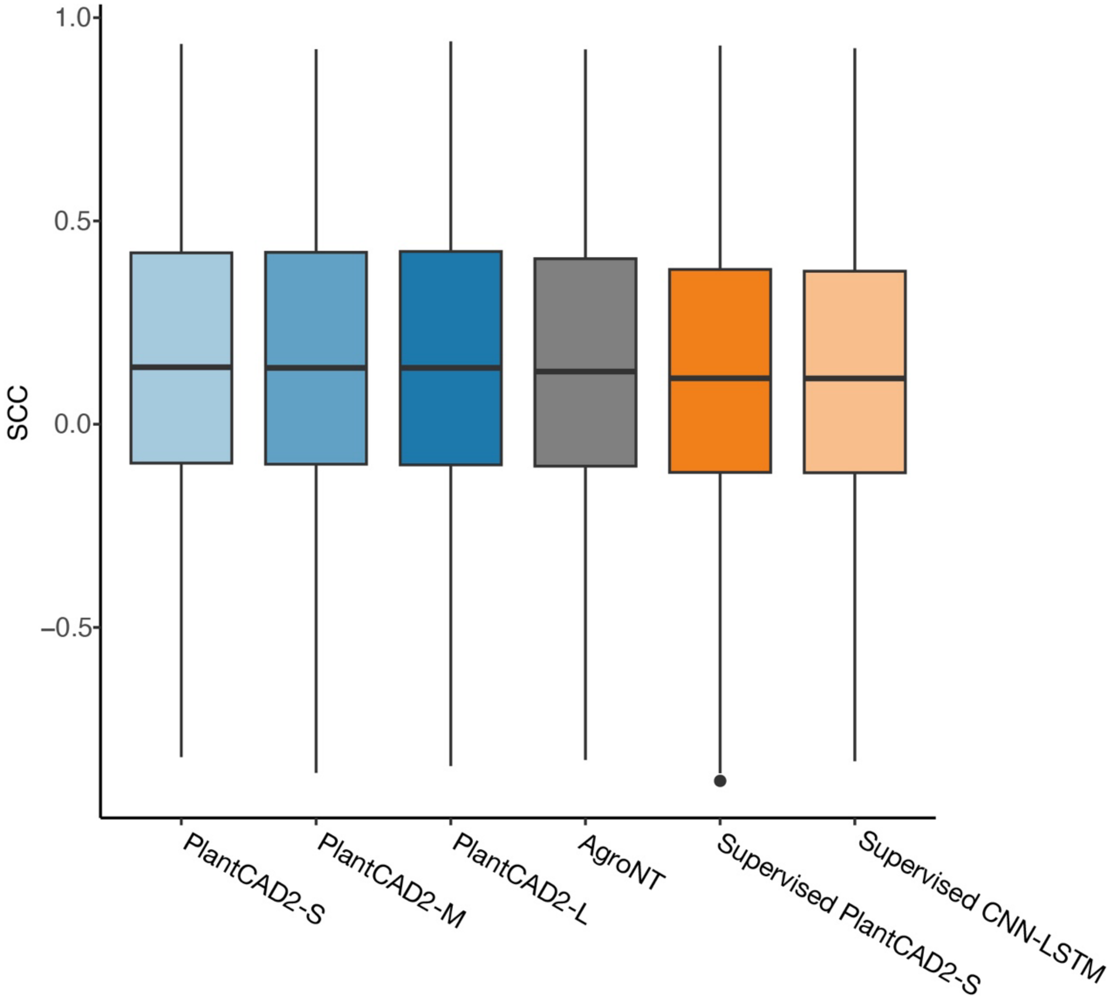


**Figure S13. Per-orthogroup expression correlation performance within the maize NAM population.** Spearman correlation coefficients (SCC) for predicting allele-specific expression differences within orthogroups.

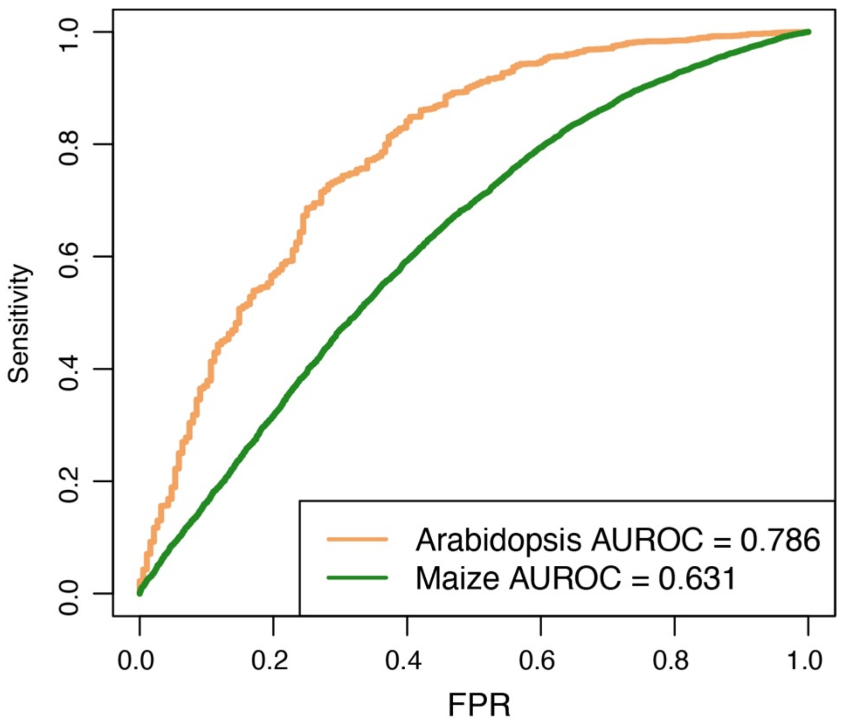


**Figure S14. Cross-species transfer performance for gene expression prediction.** ROC curves showing performance when training on Arabidopsis and testing cross-species transfer to maize.

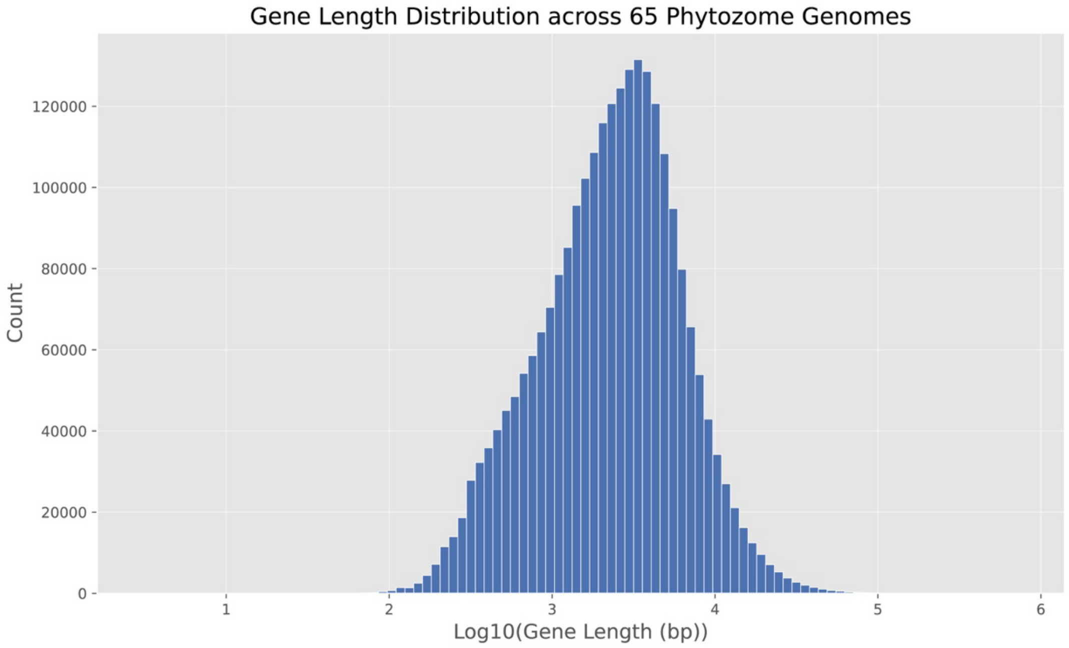


**Figure S15. Gene length distribution of 65 pretraining genomes.**

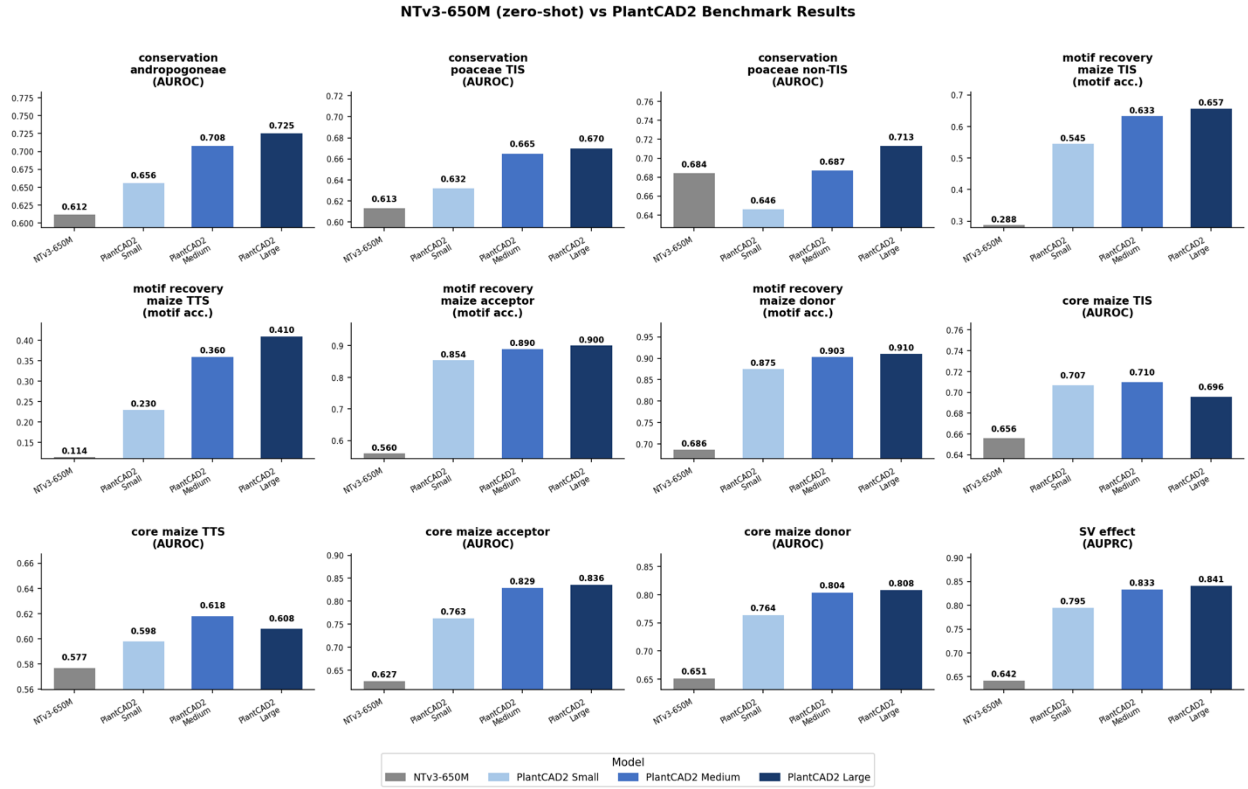


**Figure S16. Zero-shot performance comparison between PlantCAD2 and NTv3.**
